# Antimicrobial use contributes to resistance gene enrichment across cattle groups on commercial dairy farms

**DOI:** 10.64898/2026.05.22.726633

**Authors:** Andrew J. Steinberger, Colette A. Nickodem, Juliana Leite de Campos, Ashley E. Kates, Tony L. Goldberg, Nasia Safdar, Ajay K. Sethi, John M. Shutske, Pamela L. Ruegg, Garret Suen, Jessica L. Hite

## Abstract

Antimicrobial use (AMU) in agricultural systems is frequently linked to antimicrobial resistance (AMR). Yet, the scale at which AMU reshapes host-associated resistomes remains unclear. This gap arises, in part, from the scarcity of farm-level AMU data from commercial production systems. Here, we combine detailed AMU records from commercial dairy farms with metagenomic analyses of bovine fecal resistomes from calves, lactating cows, sick cows, and cull cows. At a broad level, resistome profiles were similar regardless of farm AMU. Resistance associated with historically common antibiotics, such as tetracyclines, was frequent on low- and high-AMU farms, indicating that some resistance classes are ubiquitous in dairy systems regardless of current AMU. In contrast, resistance to other drug classes varied systematically with AMU. Higher AMU was associated with increased resistance to aminoglycosides, β-lactams, and macrolides, drug classes that are critical for treating mastitis and bovine respiratory disease. Resistance gene richness and diversity were highest in calves, underscoring the importance of accounting for host traits alongside AMU when evaluating resistance patterns. Together, these findings underscore the need for detailed, farm-level AMU data to understand how management practices shape AMR and to inform strategies for sustaining the effectiveness of existing antimicrobials in agricultural and public-health contexts.

## Introduction

Understanding links between antimicrobial usage (AMU) in agricultural systems and patterns of antimicrobial resistance (AMR) has proven difficult outside of controlled laboratory settings^1–3^. Yet, resolving these relationships in real-world systems is crucial for understanding how bacteria and resistance genes evolve and persist in complex microbial communities^4–6^. Beyond basic biology, this information is essential for improving management and policy decisions^7–9^. This is particularly true in agricultural production systems, where effective antimicrobial stewardship depends on understanding how on-farm use translates into resistance outcomes^10^. Such advances could provide solutions that optimize farm productivity and animal welfare^11,12^ while also reducing public health risks posed by drug resistance^13^.

The need for detailed information on AMU and AMR in production settings is becoming increasingly urgent. Global demand for animal protein, including meat and milk, is projected to increase by approximately 60% by 2050, placing growing pressure on livestock systems to intensify production^14,15^. As livestock operations scale to meet this demand, maintaining animal health while limiting unnecessary antimicrobial exposure represents a central challenge for producers, veterinarians, and regulators alike.

In practice, the data required to link antimicrobial use and resistance in commercial production systems remain difficult to obtain^12,16^. As a result, studies examining AMU–AMR relationships in livestock systems have produced mixed findings, with some reporting strong associations and others detecting only weak or inconsistent effects^10,17^. Rather than indicating that AMU plays no role in shaping resistance, this variability likely reflects the complexity of real-world production systems. Such differences include, for example, animal physiology, management practices, and farm environments. These differences extend to regulations regarding allowable uses of antimicrobials among different animal species, age classes, and production classes based on the route of administration^8–10^. For instance, in dairy systems, systemic and oral uses directly influence the gut microbiome while local routes, such as intramammary and topical, may have minimal absorption and short-lived impacts^18,19^.

Clear associations between AMU and AMR have been demonstrated in systems where antimicrobial exposure is measured at fine resolution and animals are followed through defined life stages. For example, longitudinal studies of Danish pig production, where AMU is tracked by batch, farm, and age group, have identified drug-class–specific relationships between AMU, route of administration, and the abundance of resistance genes^17,20^. These studies underscore that AMU–AMR links can be detected when accompanying meta-data are sufficiently detailed.

In contrast, work directly linking AMU to resistance dynamics in commercial dairy systems remains limited. Comparisons between conventional and organic dairies provide valuable perspective, but are often based on isolate recovery and antimicrobial susceptibility testing. Moreover, although organic dairies in the US use little to no antimicrobials^21,22^, cattle in these systems still harbor diverse fecal resistomes, whereas cattle from conventional farms frequently carry a higher abundance of resistance genes and, in some cases, greater overall resistance diversity^23–25^. These observations suggest that resistance is already widespread in dairy environments and that differences in AMU among conventional farms may produce subtle shifts in resistance patterns, rather than dramatic changes in resistance presence.

The implications of these resistance patterns extend beyond surveillance, directly affecting the treatment of bacterial diseases that underpin dairy productivity and animal welfare. Many of the most prevalent and economically costly diseases affecting the dairy industry such as mastitis, lameness, reproductive disorders, and respiratory infections are caused by bacterial pathogens^26,27^. AMR pathogens threaten to increase disease severity, reduce treatment efficacy, and drive even greater economic losses^28^. Despite decades of research, however, we still lack the data and computational tools to conduct precision management and to optimize use of antimicrobials^29^.

Here, we advance the empirical understanding of these gaps by leveraging prior work from our group that quantified the AMU of 40 large conventional dairy farms across Wisconsin, USA^16^. From this study, we categorized farms into “low” and “high” AMU categories and then selected eight farms for more detailed longitudinal analyses of bovine fecal resistomes^16,30^. Specifically, we analyzed fecal samples collected from dairy cattle during four on-farm visits between January and October 2020. At each scheduled visit, fecal grab samples were collected from 10 randomly selected animals within each animal group (calves, lactating cows, cull cows, and sick cows), using methods as described in previous studies.^31–33^ We then pooled these individual samples per animal group at each sample timepoint for subsequent farm-level analyses. By integrating metagenomic sequencing with targeted bioinformatic and statistical analyses, we evaluated how AMU relates to the diversity, composition, and abundance of AMR genes across farms and cattle groups.

## Results

### Antimicrobial use classifications are more complex than “high” and “low”

As expected, differences in AMU practices across farms were pronounced, even after normalizing at the herd level (Fig. 1A). Overall, the range of AMU across all farms spanned approximately seven-fold in AMU daily defined dosage (DDD)/1000 herd-days. Low- and high-AMU farms differed by nearly three-fold in mean antimicrobial use DDD/1000 herd-days. High AMU farms averaging 28.6 ± 1.94 defined daily dose (DDD)/1000-herd days and low AMU farms averaging 9.76 ± 1.29 DDD/1000-herd days. Hence, although coarse, this binary categorization captured meaningful variation in overall levels of antimicrobial exposure across farms. The most commonly used antimicrobials on high AMU farms were ceftiofur, penicillin g procaine (peng), cephapirin, sulfadimethoxine, and a trimethoprim + sulfadimethoxine combination accounting for ∼85% of the total AMU across high AMU farms. On low AMU farms, the most commonly used antimicrobials were ceftiofur, cephapirin, dihydrostreptomycin, cloxacillin, and ampicillin, which accounted for ∼92% of the total AMU on low farms.

**Fig. 1.**
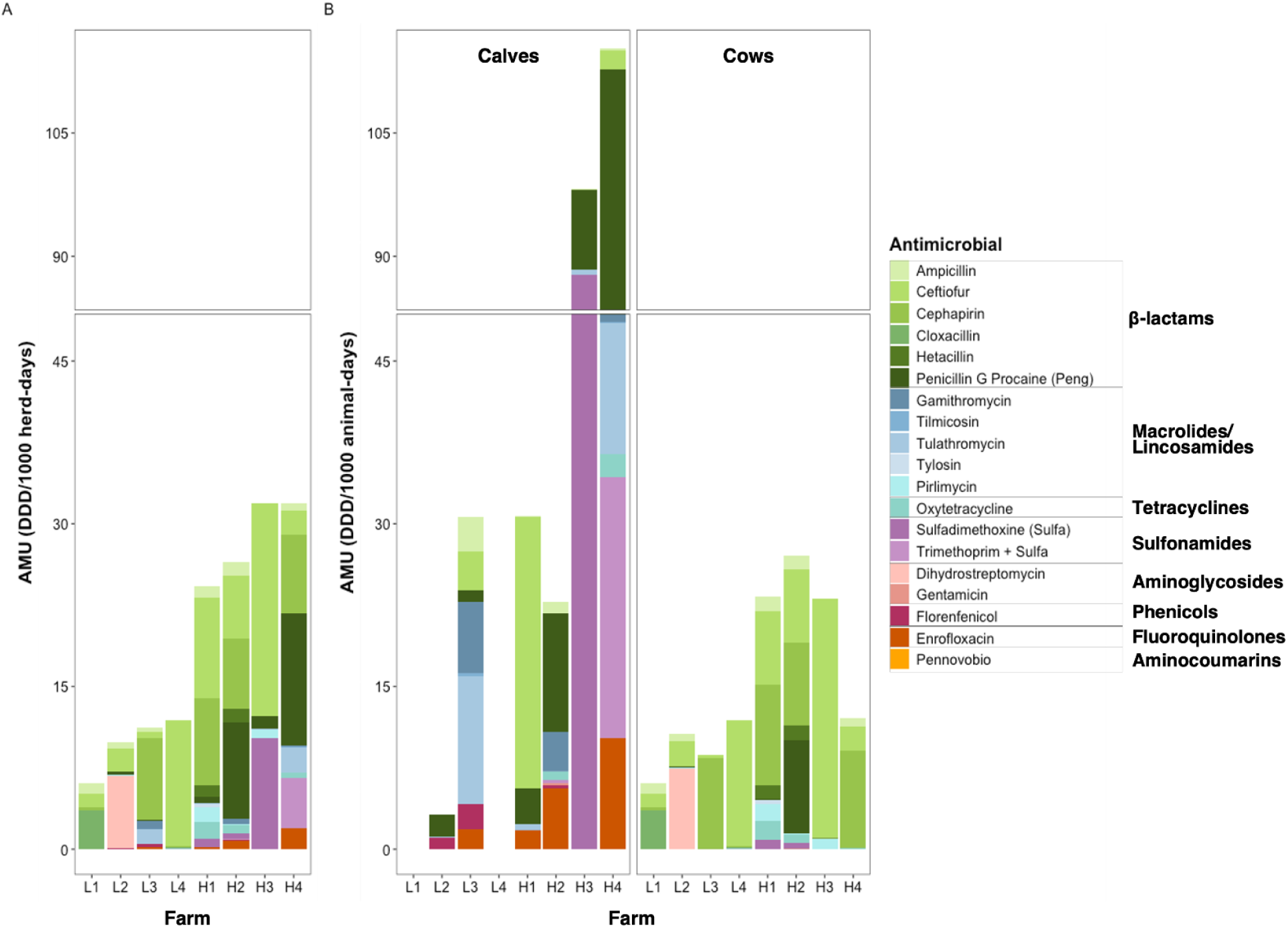
Ranking low and high antimicrobial use on dairy farms. Stacked bar plots of antimicrobial usage at dairy farms across WI, USA when normalized by the (**A**) herd or (**B**) cow age group. Usage data are summarized by the number of defined daily doses administered (DDD) and normalized by number of animals and the length of time antimicrobial use (AMU) was surveyed (365-days) multiplied by 1000. Farm labels correspond to the four low-AMU farms (L1–L4) and the four high-AMU farms (H1–H4). Farms L1 and L4 did not raise calves on-site and thus had no calf-specific AMU data reported.

Of course, farm level differences in AMU reflect herd-level variations, thus we normalized the antimicrobial data by animal group. This approach enabled a deeper investigation into the specific antimicrobials used for calves and adult cows (healthy, cull, and sick). Total calf AMU on farms H1 and H2 were more similar to those of low AMU farms L2 and L3 than the other high-use farms H3 and H4 (Fig. 1B). Use of specific antimicrobials on calves also differed between farms, with ceftiofur contributing to most treatments on farm H1, sulfadimethoxine on farm H3, peng to farm H4, H2, and L2, and Tulathromycin to farm L3. Evaluation of AMU for adult cattle (Fig. 1B) yielded similar results as herd-level AMU (Fig. 1A), except for cephapirin, which constituted the majority of AMU on H4 instead of peng when including use across all cattle (Fig. 1B). Moreover, the total AMU of farm H4 was more similar to those of low AMU farms than the other high AMU farms. These data demonstrate that binary rankings of “high” and “low” do not accurately capture the full range of potential antimicrobial exposures when data are subset to specific cattle groups for specific antimicrobial classes.

### Antimicrobial use subtly alters cattle resistomes

As a broad overview of the potential impacts of low versus high AMU, we first examined two key questions: (1) does AMU influence the number of distinct resistance genes detectable in cattle fecal resistomes (‘resistome richness’) and (2) does AMU alter which resistance classes dominate the resistome, as reflected by their relative abundances (‘resistome diversity’)?

### Antimicrobial use alters resistome richness more than diversity

Overall, higher AMU was associated with a greater number of distinct resistance genes (increased “resistome richness”) but not with a substantial shift in resistome diversity or composition. Dairy cattle resistomes from farms with high AMU were associated with increased Chao’s resistome richness (*P* = 0.03, Wilcoxon Test, Fig. 2A). In contrast, Shannon’s diversity did not differ significantly between low and high AMU farms (*P* = 0.10, Wilcoxon Test, Fig. 2B). Together, these results indicate that higher AMU was associated with a more unique repertoire of resistance genes, rather than a strong selective sweep toward specific genes.

**Fig. 2.**
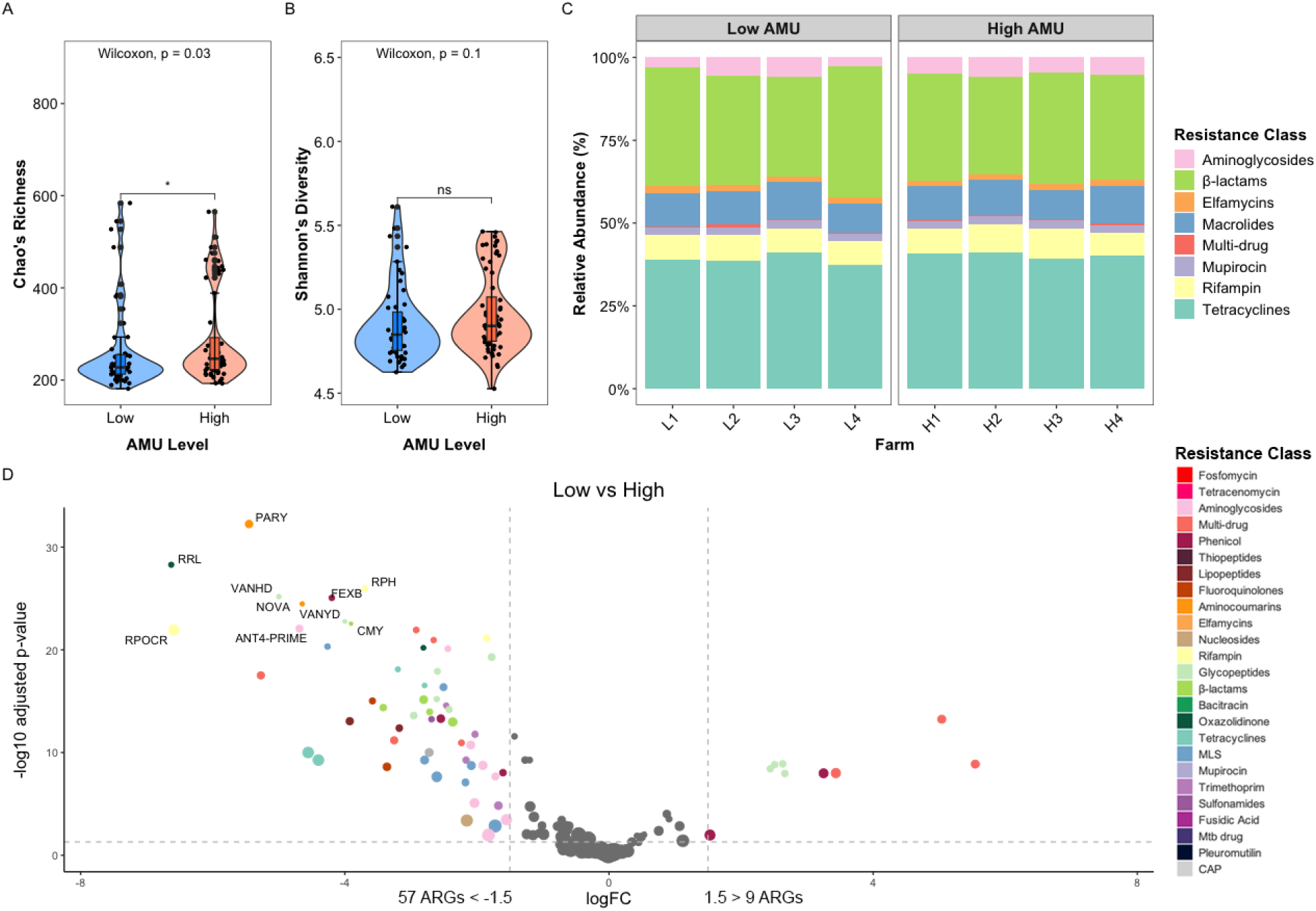
Effects of low and high antimicrobial use on the cattle resistome. Violin plots with internal boxplots of Chao’s richness (**A**) and Shannon’s diversity (**B**) for fecal samples from cattle from farms with low and high antimicrobial usage (AMU). We used Wilcoxon Rank Sum tests to determine the *P* - values of AMU level on richness and diversity, indicated by * for < 0.05. (**C**) Relative abundance of the most abundant resistance classes identified in cattle fecal resistomes across the four low-AMU farms (L1–L4) and the four high-AMU farms (H1–H4). (**D**) Resistance genes with large effect sizes (absolute log-fold change ≥ 1.5) between low- and high-AMU farms were identified as statistically significant using a zero-inflated Gaussian model (*P* < 0.05). The top 10 most statistically significant resistance genes are labeled. Compared to high-AMU farms, cattle fecal resistomes from low-AMU farms contained 57 genes with lower abundance and only 9 genes with higher abundance.

We next examined whether these differences were driven by changes in the relative abundance of genes conferring resistance to different drug classes. The relative abundance of the most prevalent resistance classes, defined as those representing > 0.5% of total gene accession counts across all samples, was similar between low and high AMU farms, with resistomes dominated by ARGs associated with resistance to β-lactams and tetracyclines (Fig. 2C). To examine associations within antimicrobial drug classes, we applied multivariate analysis and found numerous differences (PERMANOVA *P* = 0.003, *R*² = 0.086; Supplemental Table 1). Here, we included farm identity as a random effect because of the substantial variability in AMU across individual farms and used statistical tests to ensure that results were not driven by unequal dispersion among samples from the low and high AMU groups (PERMDISP *P* ≥ 0.940; Supplemental Table 1).

Tetracycline resistance was highly abundant across both low and high AMU farms (PERMANOVA *P* = 0.758, *R*² = 0.057, PERMDISP *P* = 0.810; Supplemental Table 1). Among the other most abundant antimicrobial resistance classes, resistance to aminoglycosides (*P* = 0.028, *R*² = 0.085), β-lactams (*P* = 0.015, *R*² = 0.128), and macrolides/lincosamides/streptogramin (MLS) (*P* = 0.0003, *R*² = 0.129) each varied significantly by AMU level (PERMANOVA). These variations were not driven by unequal dispersion among samples from the low and high AMU groups (all PERMDISP *P* ≥ 0.305; Supplemental Table 1) These findings are somewhat unexpected in light of the low reported use of tetracyclines across the farms included in this study (Fig. 1).

A total of 57 individual resistance genes were more abundant in cattle resistomes from high AMU farms compared to low AMU farms (|logFC| > 1.5, *P* < 0.05; zero-inflated Gaussian model; Fig. 2D). Importantly, on low AMU farms, we found a statistically significant lower abundance of the β-lactamase gene *bla_CMY_* (logFC = -3.91, *P_FDR-ad_*_j_ = 2.92 x 10^-23^), which confers resistance to 3rd-generation cephalosporins, especially in calves (logFC = -3.91, *P_FDR-ad_*_j_ = 0.02; Fig. 3A). When considering β-lactamase resistance genes in isolation, there were 2 β-lactamase resistance genes that were less abundant in calves and 6 β-lactamase resistance genes that were less abundant in adult cows on low AMU farms compared to high AMU farms (Fig. 3). Specifically, we identified a statistically significant lower abundance of β-lactamase gene *bla_CTX_* (logFC = -1.73, *P_FDR-ad_*_j_ = 1.62 x 10^-7^) in cows on low AMU farms, which confers resistance to a broad range of β-lactam antibiotics that are clinically relevant for human infections, including extended-spectrum (3rd and 4th generation) cephalosporins (Fig. 3B). Additionally, statistically significant lower abundances of *ampC* (logFC = -2.14, *P_FDR-ad_*_j_ = 1.92 x 10^-17^), which confers resistance to penicillins and most cephalosporins, were identified in adult cows on low AMU farms.

**Fig. 3.**
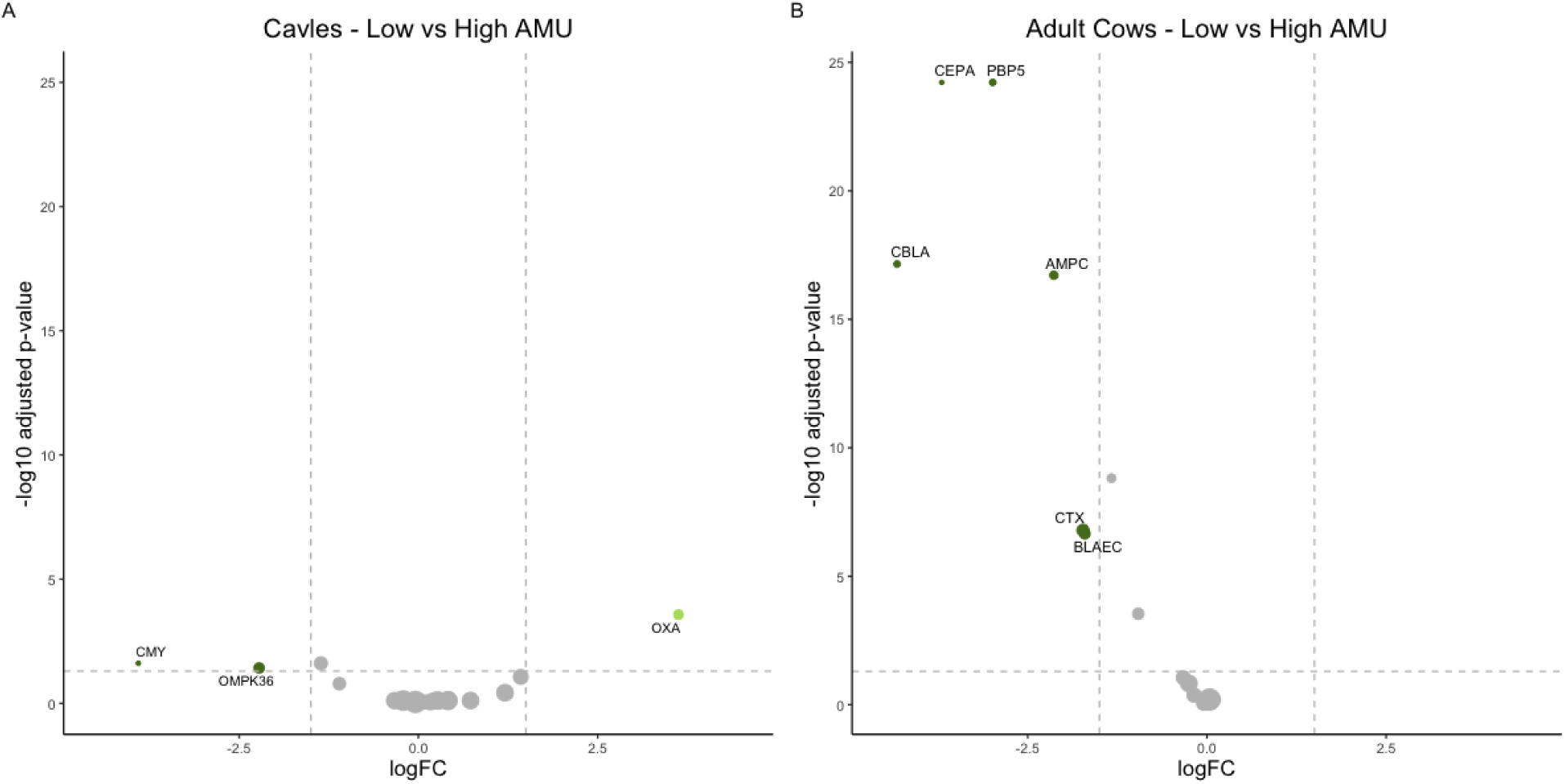
Effects of low and high antimicrobial use on the abundance of β-lactamase resistance genes in cattle. Β-lactamase resistance genes with large effect sizes (absolute log-fold change ≥ 1.5) between low- and high-AMU farms were identified as statistically significant using a zero-inflated Gaussian model (*P* < 0.05). **A)** In calf resistomes, two β-lactamase genes were significantly less abundant and one β-lactamase gene was significantly more abundant on low AMU farms relative to high AMU farms. **B)** In adult cow resitomes, six β-lactamase genes were significantly less abundant on low AMU farms relative to high AMU farms, in particular *bla_CTX_* and *ampC*, both of which are clinically relevant.

Although AMU explained a modest proportion of the variance in the composition of resistance class (8.6%), it was retained in subsequent models due to its biological relevance and its statistically significant associations with both resistome richness and overall composition. We therefore further evaluated AMU alongside additional potentially confounding variables, such as seasonality and cattle group, in downstream analyses.

### Resistome diversity shows limited seasonal variation

To evaluate temporal patterns, we examined resistome alpha and beta diversity at multiple temporal scales, including monthly and seasonal groupings of cattle across farms, as well as longitudinal within individual farms. We found no statistically significant differences in alpha diversity across months or seasons within or across farms (*P* > 0.05 for all comparisons; Supplemental Figs. 1 and 2). For beta diversity, month explained 8.8% of the variance (PERMANOVA *P* = 0.106; PERMDISP *P* = 0.500), season explained 8.2% (PERMANOVA *P* = 0.103; PERMDISP *P* = 0.380), and longitudinal sampling at individual farms explained 8.5% (PERMANOVA *P* = 0.051; PERMDISP *P* = 0.555). Although longitudinal sampling approached significance, it was conducted at inconsistent intervals and none of the temporal variables were consistently associated with differences in resistome structure across alpha or beta diversity metrics. Consequently, these variables were not retained in subsequent analyses.

### Resistome alpha diversity differs by cattle groups

Both Chao’s richness (Kruskal–Wallis *P_FDR-ad_*_j_ = 1.2 × 10⁻¹²) and Shannon’s diversity (Kruskal–Wallis *P_FDR-ad_*_j_ = 9.6 × 10⁻¹³) differed across cattle groups, including calves, cull cows, sick cows, and healthy lactating cows. Calves exhibited higher richness and diversity than all other groups (Kruskal–Wallis *P_FDR-ad_*_j_ < 0.001), and sick cows also had higher fecal resistome diversity than cull cows (Kruskal–Wallis *P_FDR-ad_*_j_ < 0.05; Supplemental Fig. 3; Supplemental Table 2). These animal-group specific patterns were consistent when analyses were stratified by AMU level, with statistically significant differences in richness and diversity observed at both high (Kruskal–Wallis *P_FDR-ad_*_j_ = 1.3 × 10⁻⁸ and 3.4 × 10⁻⁸, respectively; Fig. 4A) and low (Kruskal–Wallis *P_FDR-ad_*_j_ = 4.5 × 10⁻⁵ and 1.6 × 10⁻⁵, respectively; Fig. 4B) AMU farms, although specific pairwise comparisons varied.

**Fig 4.**
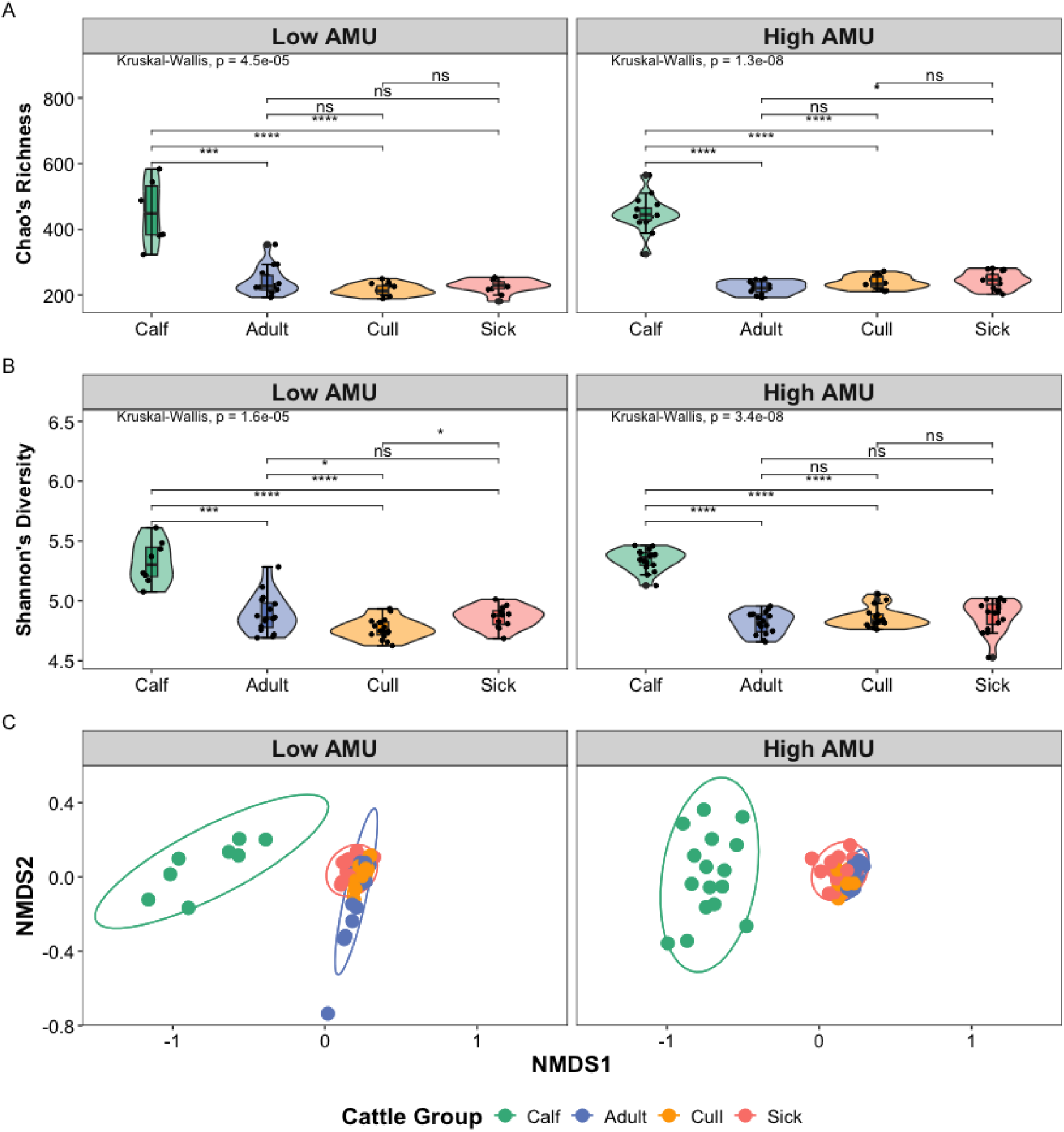
Comparison of resistome diversity between cattle groups. Violin plots with internal boxplots of (**A**) Chao’s richness and (**B**) Shannon’s diversity for fecal samples from each of the 4 cattle groups from low and high antimicrobial use (AMU) farms. Differences in diversity among cattle groups were first evaluated using a Kruskal–Wallis test. When significant, pairwise differences between cattle groups were assessed by Wilcoxon rank-sum tests using FDR-adjusted P-values, with < 0.05 indicated by *, < 0.001 by ***, and < 0.0001 by ****. (**C**) Non-metric multidimensional scaling (NMDS) ordination for Bray-Curtis dissimilarity analyses of fecal resistomes across cattle group and antimicrobial usage (AMU). Ellipses represent standard errors around the centroid for each group (Stress = 0.058, Non-metric fit R^2^ = 0.997; See Supplemental Fig. 4).

### Resistance class composition differs across cattle group

The composition of genes conferring resistance to different drug classes also differed among cattle groups, driven largely by pronounced variability in calves. While differences in resistance were maintained across AMU levels for all other cattle groups, differences between adult and cull cattle emerged only under low antimicrobial use conditions. We found statistically significant differences in specific resistance classes between cattle groups (PERMANOVA *P* < 0.0001, Supplemental Table 3). However, within groups dispersion of Bray-Curtis’s dissimilarities were heterogeneous (PERMDISP *P* = 0.022) due to dispersion in calves (Fig. 4C). Pairwise comparisons, excluding calves, revealed that all cattle groups (Adult vs Sick and Cull vs Sick) varied significantly in the abundance of genes conferring resistance to different drug classes (PERMANOVA *P* <0.001; PERMDISP *P* = 0.2) except for adult and cull cattle (*P* = 0.118), which remained consistent for high AMU (*P* = 0.079), but not low AMU (*P* = 0.017) (Supplemental Table 4).

For calf samples, tetracycline resistance genes were the most abundant (48.8±3.9%), followed by β-lactams (16.8±4.0%), MLS (macrolide, lincosamide, and streptogramin) (11.2±1.7%), and aminoglycosides (1.1±0.4%) (Supplemental Fig. 5). In contrast, the resistome of adult cattle was dominated by β-lactam (36.8±5.0%) and tetracycline (36.5±4.3%) resistance, with lower relative abundances of MLS (9.7±1.3%), rifampin (8.5±2.3%), and aminoglycosides (3.0±0.57%) (Fig. 5A). All of the most abundant resistance classes (aminoglycosides, β-lactams, MLS, and tetracyclines) varied significantly by cattle group (PERMANOVA *P* < 0.0001; PERMDISP *P* > 0.05) (Fig. 5A). Elfamycin, mupirocin, and rifampin also varied significantly by cattle group (PERMANOVA *P* < 0.0001), but dispersion of these were non homogenous (PERMDISP *P* < 0.05) (Supplementary Table 3).

**Fig 5.**
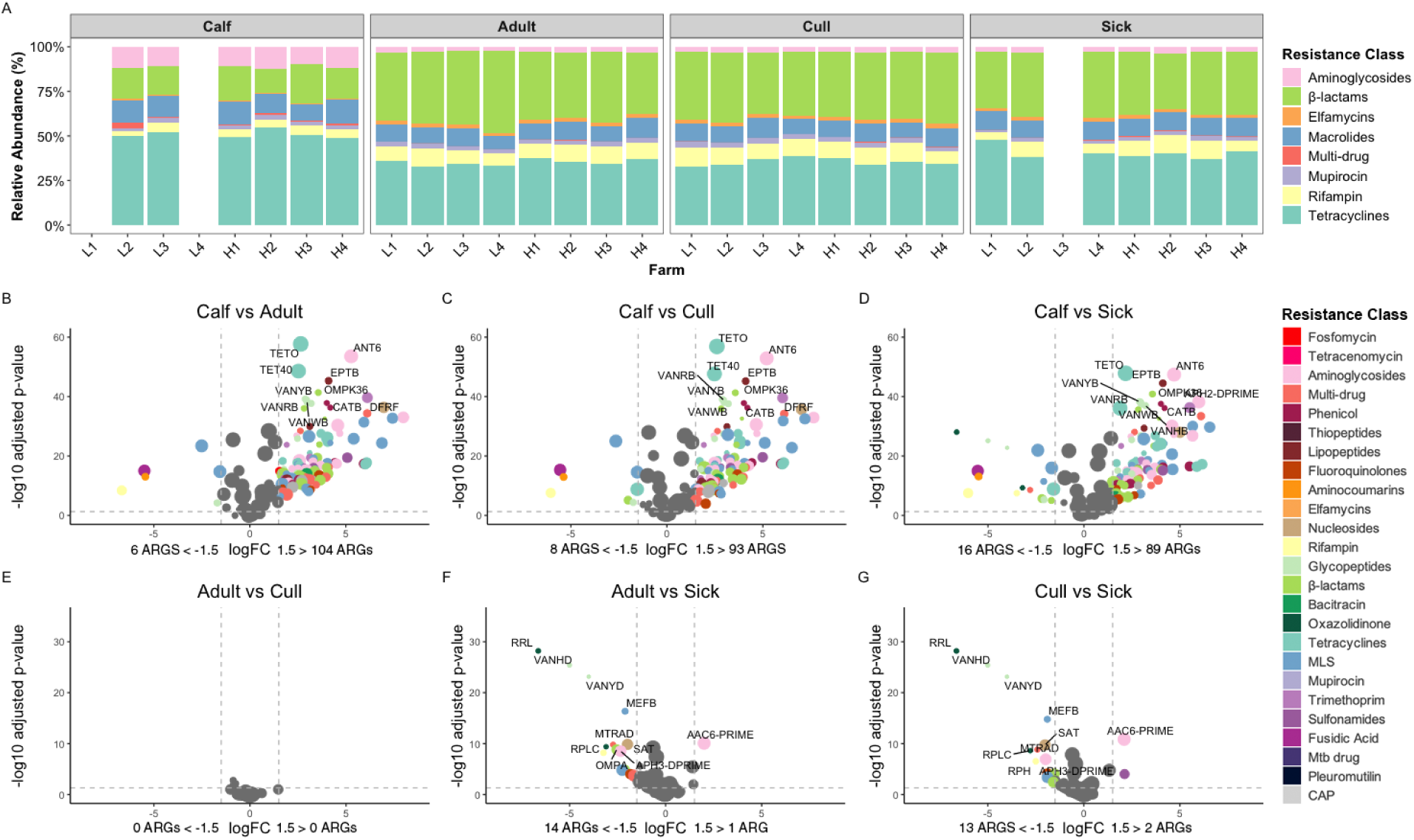
Resistance patterns differed by drug-class across cattle groups. (**A**) Relative abundances of genes conferring resistance to specific antimicrobial classes across farms and cattle groups. We subset data to include resistance classes representing > 0.5% of total gene accession counts across all samples. Farms L1 and L4 did not raise calves and there were no sick animals on farm L3. (**B-G**) Differentially abundant antimicrobial resistance genes amongst pairwise comparisons between cattle groups (*P* < 0.05), with logFC ≤ 1.5 on the left and logFC ≥ 1.5 on the right. Vertical gray dotted lines represent |logFC| cutoff. Dot size indicates the magnitude of the logFC and color indicates the antimicrobial class. For plots B-D, the top 10 most statistically significant differentially abundant genes are labelled and for plots E-G all statistically significant differentially abundant genes are labelled. MLS: macrolides/lincosamide/streptogramin, Mtb: *Mycobacterium tuberculosis*, CAP: cationic antimicrobial peptide.

To identify pairwise differential abundances (DA) of resistance classes between cattle groups, we accounted for AMU and farm variability while using mixed directional false discovery rate (mdfdr) adjusted *p*-values (*q* < 0.05). Here, we only present robust DA results that passed our sensitivity analysis. Relative to adults, calves carried higher abundances of aminoglycoside (1.05 logFC), bacitracin (0.96 logFC), cationic antimicrobial peptides (1.11 logFC), fosfomycin (1.41 logFC), and multi-drug resistance (1.67 logFC) (Supplemental Fig. 6). Of note, calves consistently exhibited abundances of β-lactams, elfamycins, and rifampin that were ∼1 to 1.5 log fold-change lower compared to other cattle groups (Supplemental Fig. 6). Tetracycline resistance was only differentially abundant between calves and cull cows (logFC = 0.64, *q* = 0.013). These results may have arisen, in part, from intrinsic resistance or standing genetic variation in the farm environment, as elfamycins and rifampin are never used in dairy cattle and tetracycline usage was notably low in the farms included in this study.

### Resistance genes and mechanisms differ systematically across cattle group

Next, we examined whether these patterns were driven by differences in the underlying resistance genes and their specific mechanisms of resistance. To identify shifts in the relative abundance of resistant genes between cattle groups, we used zero-inflated gaussian models to identify statistically significant changes (|logFC| > 1.5, *P* < 0.05). Calf resistomes were enriched in several ARGs relative to adult (n = 104 |logFC| > 1.5; Fig. 5B), cull (n = 93 |logFC| > 1.5; Fig. 5C), and sick animals (n = 89 |logFC| > 1.5; Fig. 5D). We found that tetracycline resistant genes, *tetO* and *tet40*; aminoglycoside resistant genes, *ant(6)* and *aph(2”)*; β-lactamase carbapenem resistant gene, *ompK36*; vancomycin resistant genes, part of the *vanB* resistance cluster (*van*YB, *van*RB, and *van*WB); and chloramphenicol resistant gene, *catB,* were among the most differentially abundant genes in calves. Resistomes from sick animals were also enriched with a smaller set of ARGs compared to adult (n = 14 |logFC| > 1.5; Fig. 5F) and cull (n = 13 |logFC| > 1.5; Fig. 5G) animals. We identified no major differences in the abundance of resistance genes between adult and cull animals (Fig. 5E).

To identify variation in specific resistance mechanisms between cattle groups, we investigated resistance mechanism composition and relative abundance. In calf samples, tetracycline ribosomal protection proteins (46.1±3.6%) were by far the most abundant ARG mechanistic class, followed by class A β-lactamases (15.7±3.8%). Additionally, 23S rRNA methyltransferases (5.1±1.3%), rifampin-resistant RpoB (4.3 1.2%), both aminoglycoside O-nucluotidyltransferases (4.9±05.2%) and aminoglycoside O-phosphotransferases (4.2±06.4%), and macrolide-resistant 23S rRNA mutation (3.1±6.9%) were also highly abundant (Supplemental Fig. 7). In adult cattle the resistome was dominated by class A β-lactamases (36.6±5.1%), tetracycline ribosomal protection proteins (35.5±4.4%), and rifampin resistant RpoB (8.5±2.3%). These results matched the overall trends in the relative abundance of genes conferring resistance to specific drug classes described above (Fig. 5A).

Given the complexity of these results, we next examined the resistance gene and mechanism compositions of the top 4 most abundant resistance classes only. Consistent with earlier patterns, calves exhibited a disproportionately large repertoire of unique resistance genes relative to other cattle groups (Fig. 6). While adult cows showed higher overall abundance of β-lactam resistance, calves contributed a greater diversity of unique β-lactam resistance genes, many of which are clinically relevant (Fig. 6B). Similar trends were observed across aminoglycoside, MLS, and tetracycline resistance genes (Fig. 6A, 5C, 5D, and Supplemental Figs. 8–11).

**Fig 6.**
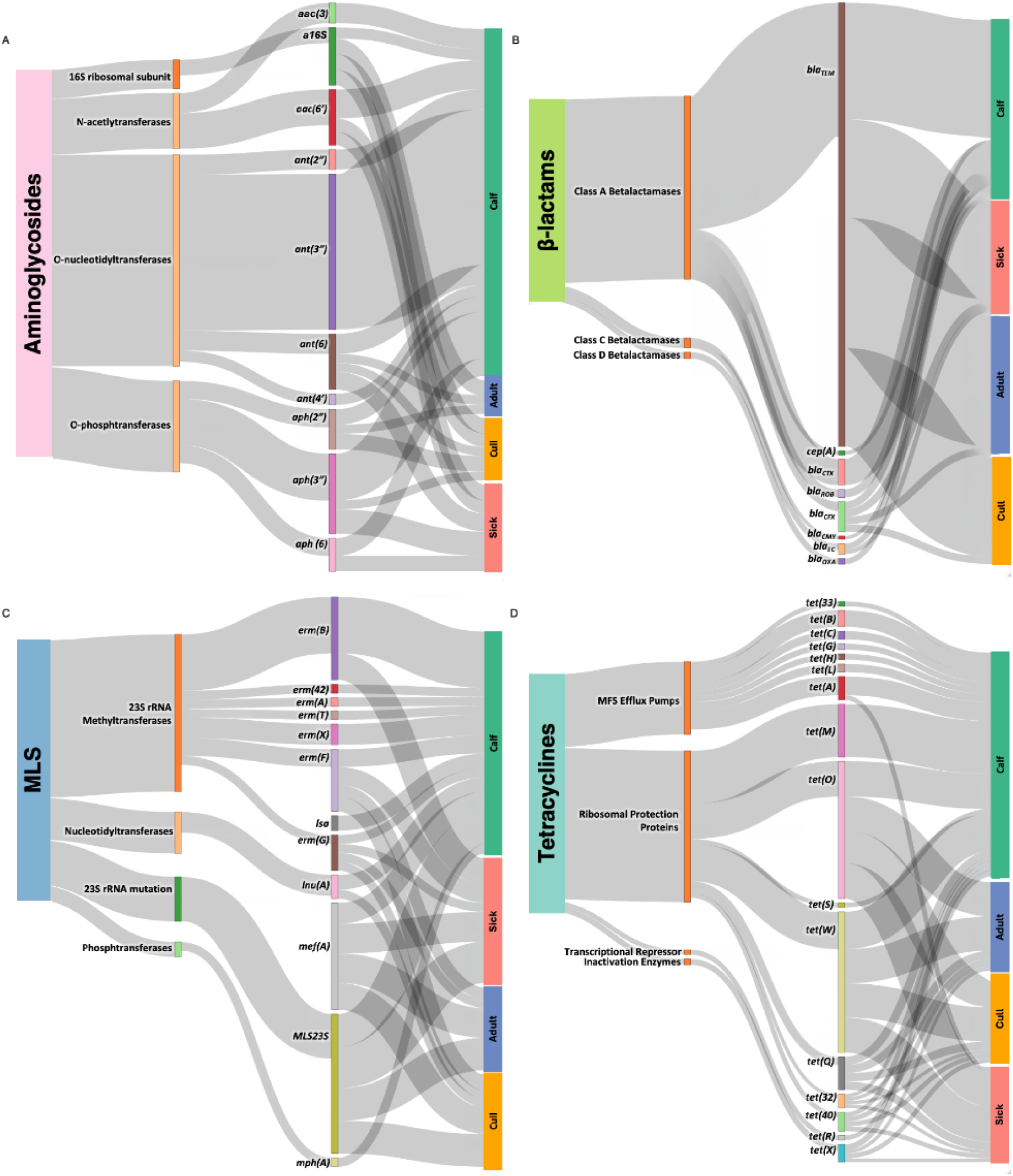
Sankey plots of the top 4 most abundant antimicrobial resistance classes: (A) aminoglycosides, (B) β-lactams, (C) macrolides/lincosamides/streptogramin (MLS), and (D) tetracyclines. These plots show the proportion of genes attributed to each resistance mechanism and the proportion of genes identified in each cattle group. In all cases, the highest proportion is attributed to calves and in most cases, there is a higher diversity of unique resistance genes attributed to calves.

### Host characteristics shape AMU–resistome relationships

Finally, we subset the data into calves and adult cattle to evaluate AMU (animals/ defined daily dose (DDD)) as a continuous variable rather than the binary variable used above. Within calves, resistome alpha diversity did not correlate with calf-specific AMU (calf/DDD), using either Chao’s richness (Spearman’s *r* = 0.058, *P* = 0.787) or Shannon’s diversity (Spearman’s *r* = 0.148, *P* = 0.490; Supplemental Table 5). In contrast, the composition of calf resistomes (beta diversity) differed across AMU (calf/DDD) gradients (PERMANOVA *P* = 0.01, *R*^2^ = 39.1%, PERMDISP *P* = 0.527). These results suggest that AMU levels alter the structure, but not the overall richness, of the calf resistome.

Among adult cattle, increasing AMU (cow/DDD) was positively correlated with Chao’s richness (Spearman’s *r* = 0.242, *P* = 0.020), but had no effect on Shannon’s diversity (Spearman’s *r* = 0.142, *P* = 0.176). Resistome compositions also varied with AMU (cows/DDD) (PERMANOVA *P* = 0.04, *R*^2^ =11.9%, PERMDISP *P* = 0.23). As expected, these analyses using quantitative metrics of AMU (cows/DDD) (i.e., as a continuous variable) captured substantially more variation in diversity and composition of resistomes than binary AMU categorization. While these findings are biologically intuitive, such relationships are rarely examined in livestock systems due to the difficulty of collecting high-resolution, quantitative antimicrobial use data.

To assess whether the relationship between AMU and resistome diversity varied by health status, we evaluated each subgroup of adult cows individually (healthy, cull, and sick cows). However, viewed in this manner, the impact of AMU (cows/DDD) on resistome alpha diversity did not differ across subgroups. Among healthy cows, neither ARG richness (Chao’s richness: *r* = 0.018, *P* = 0.920) nor diversity (Shannon’s (diversity: *r* = 0.043, *P* = 0.816) varied with AMU. In contrast, cull cows exhibited a strong positive relationship between AMU and ARG richness (Chao’s richness: Spearman’s *r* = 0.406, *P* = 0.021) and Shannon’s diversity (Spearman’s *r* = 0.388, *P* = 0.028). Among sick cows, ARG richness was marginally correlated with AMU (*r* = 0.369, *P* = 0.053), whereas Shannon’s diversity remained unchanged (*r* = 0.093, *P* = 0.634).

Despite these minimal subgroup-specific differences in alpha diversity, resistome composition varied significantly with AMU across all adult cow subgroups (Bray–Curtis PERMANOVA). AMU (animals/DDD) explained substantial variation in resistome composition among healthy cows (*R²* = 35.4%, *P* = 0.03), cull cows (*R²* = 38.9%, *P* = 0.005), and sick cows (*R²* = 37.9%, *P* = 0.008), with no evidence of differential dispersion within groups (all PERMDISP *P* > 0.08).

## Discussion

Here, we find that antimicrobial use (AMU) shapes resistance patterns in commercial dairy systems in ways that are obscured when data are aggregated across animals. Specifically, calves from farms with high AMU harbored higher abundances and more diverse β-lactam resistance genes, many of which are clinically relevant^34,35^. Together, drug usage and animal group explained approximately 86% of the variation in resistome patterns. However, when using the standard approach of aggregating across animal groups, resistome class composition appeared largely similar and standard “low” and “high” AMU explained only 8% of the observed variation.

Together, these results help resolve a recurrent puzzle: AMU–AMR relationships in livestock systems have produced mixed findings, with some reporting strong associations and others detecting only weak or inconsistent effects^10,17^. For example, *Doster et al.* (2022) reported that AMU effects on beef cattle fecal resistomes were sometimes statistically detectable, but explained less than 1% of total variation, leading to the conclusion that AMU is not a strong driver of resistance emergence in that system^36^. Our results suggest that previous studies typically investigate AMU-AMR links on the wrong scale. Because detailed, farm-level AMU records from commercial dairies, paired with resistome data across cattle groups, are exceptionally difficult to assemble^12,16^, AMU effects are often evaluated using coarse summaries. Under those conditions, averaging across different animal groups masks AMU signals beneath larger differences in age, health status, and management practices^17,20^. Given the substantial differences in both the type and amount AMU across farms, the mixed results from previous studies are not surprising.

At the herd level, higher AMU was associated with a greater number of distinct resistance genes, indicating that cattle on high-AMU farms carried a broader set of resistance determinants. This increase in richness did not coincide with a change in overall resistome diversity, meaning that the same resistance types continued to dominate across farms regardless of AMU level. In both low- and high-AMU systems, resistance was largely structured around tetracycline- and β-lactam-associated genes. We also found resistance to elfamycins and rifampin, which are never used in dairy production systems^7,9,16^. The persistence of these resistance classes across farms indicates that contemporary AMU shapes the breadth of resistance determinants present but does not substantially alter the core structure of the resistome.

For tetracyclines in particular, we observed high resistance gene abundance despite notably low reported use on the focal farms^16^. This pattern is consistent with the broader resistome concept, which encompasses resistance genes harbored by environmental and commensal bacteria, many of which predate modern clinical and agricultural antibiotic use^4,13^. Tetracyclines also have a long history of widespread use in both human medicine and livestock production, including dairy systems, contributing to their pervasive presence across microbial communities^37,38^. Consequently, the observed prevalence of tetracycline resistance likely reflects multiple, non-mutually exclusive mechanisms, including intrinsic resistance within farm environments, legacy effects of historical AMU, co-selection driven by other antimicrobials, metals, or biocides, and persistence within shared reservoirs such as soils, manure, and water^39,40^. Continued environmental exposure and transfer across animal cohorts may further maintain these resistance determinants even in the absence of strong contemporary selection pressure.

Crucially, this similarity in dominant resistance types across farms does not imply that resistomes were unaffected by antimicrobial use. Instead of “rebuilding” the resistome, higher AMU was associated with increases in *particular* resistance genes and resistance mechanisms within an already shared background. This pattern was most evident at finer gene-level resolution, where dozens of individual resistance genes were more abundant on high-AMU farms, including clinically relevant β-lactamase genes such as *bla*_CMY_ and *bla_CTX_*^34,35^. Together, these results suggest that AMU influences *how much* resistance is present and *which genes increase*, without fundamentally changing the overall genetic architecture of the resistome.

Two related features of commercial dairy systems contribute to the patterns found here. First, “total AMU” can conceal major differences in which drugs are used, how intensively they are used, routes of administration, and which animals receive them^8–10^. Second, farms differ markedly in the age structure and health status of animals being sampled, and those differences strongly shape microbial communities and resistance gene profiles. Consistent with these considerations, our data show that cattle groups, accounting for farm-to-farm differences, explained more variation in resistome patterns than herd-level AMU categories alone, even when AMU was associated with detectable shifts in composition.

More specifically, when we examined AMU on a finer scale and stratified our data by age and health status, additional trends appeared that would otherwise have remained obscure. This underscores that AMU is a meaningful contributor to resistance dynamics on dairy farms, but that its signal is easiest to detect when the analysis aligns with how antimicrobial exposure and selection actually occur in practice^1,3,10^. While this is an intuitive result, data on specific types and amounts of antimicrobial use on commercial production systems remain exceptionally difficult to obtain, such that no studies prior to ours have achieved the level of analytical resolution presented here.

Our results carry important implications for how AMU–AMR relationships are measured and interpreted. For instance, subsetting AMU to calves versus adult cows revealed that the original “high” and “low” farm classifications did not always reflect antimicrobial exposure for a given cattle group. Accounting for both cattle group and AMU revealed that calves and sick cows had distinct resistomes relative to healthy and cull cows, including higher richness and diversity and clear differences in composition. These findings align with prior work showing that calves often carry more diverse and abundant resistomes than adult cattle^41–43^. This finding is notable given that calves have relatively higher exposure to antimicrobials, though over a shorter time frame. Typically, calves are placed in a preweaned group for 45-60 days, after this timeframe they are rarely ill and rarely receive antimicrobials until they calve and enter the milking herd around 22-24 months of age.

In addition to differences in drug approval by physiological class, the route of antimicrobial administration varies substantially between calves and adult cows and likely influences observed resistance patterns^18,19^. Preweaned calves are treated exclusively via systemic or oral routes, whereas adult cows receive antimicrobials primarily through intramammary (IMM) administration^18,19^, with more limited systemic use and no oral exposure^11,12,16^. Several commonly used antimicrobials differ in their available routes: for example, ceftiofur and penicillin can be administered both systemically and intramammarily, whereas cephapirin, cloxacillin, and dihydrostreptomycin are restricted to IMM use^8–10^.

In contrast, sulfadimethoxine and trimethoprim-sulfonamide combinations are administered systemically, and ampicillin is available via both routes. These distinctions are potentially important when comparing resistance patterns across age classes/animal groups, as systemic and oral administration are more likely to exert selection pressure across broader commensal microbial communities in the digestive system, while IMM treatments are limited to more localized selection of pathogens residing within the mammary gland^18,19^. Consequently, differences in both antimicrobial availability and route of exposure between calves and cows may contribute to distinct resistome signatures, even within the same farm environment.

Overall, the elevated and distinct resistomes observed in calves and sick cows suggest that targeted, group-specific antimicrobial stewardship may be more effective than blanket farm-level policies alone. This finding has direct implications for future intervention efforts and warrants deeper investigation. Notably, relatively few studies have focused specifically on the resistomes of sick cows. The limited work on sick or culled cows indicates that antimicrobial treatment can lead to both short- and longer-term increases in resistance genes, including increases following cephalosporin treatment (e.g., intramammary ceftiofur) and enrichment of β-lactamase resistance genes after 3rd-generation cephalosporin use^44,45^. The elevated resistome diversity observed in sick cows in our study is consistent with these patterns, particularly because each sample represents a composite from multiple animals with different drugs, doses, and stages of treatment at the time of sampling.

Our work is not without limitations. First, we acknowledge that the majority of AMU in dairy cattle is for treatment of mastitis, with most drugs administered via intramammary (IMM) infusion and a smaller proportion given systemically, but not orally. As a result, much of the AMU in adult cows is delivered locally to the mammary gland, an organ that is not generally considered to harbor a stable, commensal microbiota comparable to that of the gastrointestinal tract. While some antimicrobials administered intramammarily or systemically (e.g., ceftiofur, penicillin, ampicillin) are absorbed and can transiently affect the gut microbiota, prior work suggests these effects are relatively short-lived. Consistent with this interpretation, relatively little resistance development has been documented among mastitis pathogens, potentially because only the teat canal, rather than the mammary parenchyma, harbors a commensal microbiota, resulting in limited selection pressure for resistance.

In contrast, the gastrointestinal tract contains a dense and metabolically active commensal microbiota that is highly responsive to antimicrobial exposure, including indirect effects arising from IMM or systemic antimicrobial use ^44,46^. For this reason, we focused on fecal samples, where the downstream impacts of AMU on resistance gene diversity and composition are most likely to be detectable. Perhaps more importantly, we also focused on fecal samples because they can serve as a conduit for transmitting ARGs through direct (e.g., cow-to-cow) and indirect (environment-to-human) pathways. Our future studies are investigating these dynamics to identify critical transmission points across these agroecosystems using quantitative risk analysis (Nickodem et al., in prep). Moreover, the associations between AMU and resistance patterns observed here are likely conservative, and would be expected to be stronger in production systems where antimicrobial administration occurs primarily via the oral route.

Additionally, our AMU data were collected 2.5 years prior to animal sampling on farms due, in part, to delays resulting from the SARS-CoV2 pandemic (see Supplementary Information). Although farm managers reported no substantial changes in AMU practices during this period, our analyses therefore reflect relationships with historical farm-level AMU, rather than antimicrobial exposures immediately preceding sampling. Linking resistome measurements with contemporaneous AMU data will be an important next step for refining temporal associations between use and resistance. Third, we were unable to explicitly quantify several farm-level factors that likely influence resistome composition, including diet, housing environment, bedding, and other management practices. These factors vary widely across commercial dairies and may interact with AMU to shape resistance patterns. Expanding this work to include more detailed metadata and a larger number of farms will be critical for disentangling the relative contributions of antimicrobial and non-antimicrobial drivers.

An additional limitation was the absence of calf samples from two low-AMU farms, which constrained direct comparisons of calf resistomes across the full spectrum of herd-level AMU. However, evaluating AMU as a continuous variable and conducting analyses within cattle groups likely mitigated some of this limitation. Finally, our results are based on relative resistance gene abundances, rather than absolute resistance burdens. While metagenomic profiling provides robust insight into resistome structure and composition, future studies that integrate metagenomics with absolute quantification approaches (e.g., qPCR or ddPCR) will be essential for assessing how AMU influences the total load of resistance genes carried by cattle ^47,48^.

This study provides a rare, field-based view of how AMU relates to resistome patterns in large commercial dairy systems. Given the challenges of linking detailed AMU data with resistance measurements in commercial settings^12,16^, this study offers an important step forward, while also revealing areas where additional data are needed.

### Conclusions

By leveraging detailed antimicrobial use (AMU) records linked to metagenomic resistomes across cattle groups, our study demonstrates that antimicrobial selection is detectable in commercial systems and that cattle group–specific analyses are essential for interpreting resistance dynamics^17,20^. Looking forward, continued progress will require integrating finer-scale AMU data with approaches that resolve the taxonomic and genomic context of resistance genes. Metagenome assembly and metagenome-assembled genomes will be critical for identifying the microbial taxa and mobile elements carrying ARGs, while absolute quantification approaches will help distinguish changes in resistome composition from changes in overall resistance burden. Finally, extending this work to evaluate how antimicrobial resistance associated with dairy cattle may disseminate beyond farms, including potential transfer to farm workers, will be important for understanding the broader implications of resistance in dairy production systems ^1–4^.

## Methods

### Farm selection

Farms enrolled in this study were selected from a set of 40 commercial dairy farms identified during previous research conducted by our group to quantify the antimicrobial use (AMU) of conventional dairy farms in Wisconsin^16^. Briefly, the study collected and summarized 1 year (Fall 2016 - 2017) of antimicrobial usage data for farms meeting the following enrollment criteria: 1) housed ≥ 250 lactating dairy cows, 2) use of antimicrobials to treat or prevent at least one disease event in the previous year, and 3) kept digital records of their antimicrobial treatments. AMU was quantified using 2 methods: 1) number of animal-defined daily doses administered, normalized per 1,000 animal-days (DDD), and 2) mg of antimicrobial administered per kg of animal weight (mg/kg). Using the relative distribution of the DDD metric, we identified 4 “high” and 4 “low” AMU farms from the original 40 commercial farms and contacted them via letter and/or phone to re-enroll them for bovine fecal sampling.

### Sample collection and pooling

Four bovine fecal sampling events were performed between January and October of 2020 using a schedule and sampling method adapted from prior studies^31–33^. During each visit, samples were collected from 10 individual animals, if available, from 4 different cattle groups: healthy lactating cows, cows to be culled, cows in a designated sick pen, and pre-weaned heifer calves.

Individual animals were selected via systematic random sampling. To do so, the total number of animals in the group to be sampled (*N*) were divided by the number of samples to be collected from that group (*n*) to determine a sampling interval *k* (*N*/*n* = *k*). Then every *k*^th^ animal in that group, equally distributed across pens and barns, was sampled. During each visit, cattle groups were sampled in order from calves, healthy adult and cull cow, and lastly sick cows. No effort was made to avoid collecting samples from the same individual animal in a herd during subsequent visits or to avoid sick calves when conducting systematic random sampling.

During this study, animals were sampled (n = 1,145) during 32 visits to the 8 enrolled farms, resulting in 116 composite fecal samples. Of these composites, 24 were from calves, 32 from healthy lactating cows, 32 from cull cows, and 28 from sick cows. Broken down by AMU group, low AMU farms generated 52 samples (calf: 8, healthy adult: 16, cull: 16, sick: 12) and high AMU farms generated 64 samples (calf: 16, healthy adult: 16, cull: 16, sick: 16). Discrepancies in sample counts result from two low AMU farms (L1 and L4) not rearing calves on-site and one low AMU farm (L3) not maintaining a sick pen to designate cows under antimicrobial treatment, preventing sample collection. Additionally, the sick pen of farm L4 contained fewer than 10 cows each visit so the composite sick samples for these visits comprised 3-9 individual fecal samples rather than 10 as in all other samples.

Fecal samples (∼40 g) were collected from each animal’s rectum using either shoulder length gloves (Nasco, Fort Atkinson, WI) for cows or nitrile gloves (Ansell, Iselin, NJ) for pre-weaned calves. Samples were placed into sterile 50 mL conical tubes (Falcon, Fisher Bioscience, Waltham MA) and immediately stored on wet ice before being transported to the laboratory and stored at -80 °C. To avoid cross contamination between farms, supplies stocks (glove boxes, falcon tubes, etc.) were kept separate for each farm, all clothing was laundered, and boots and equipment were washed and sanitized after each visit.

In the laboratory, each of the individual fecal samples collected from each cattle group per visit were thawed and combined at equal mass (∼10g where available) into sterile Whirl-Pak bags (Nasco) and homogenized by hand squeezing (1-2 min) until visibly mixed. A total of ∼2 g of the resultant composite sample was then transferred into sterile 2 mL screw-cap tubes (VWR / Avantor, Radnor, PA), assigned randomized labels, and stored at -80 °C until DNA extraction.

### DNA extraction and shotgun sequencing

DNA was extracted directly from 222 – 253 mg of composite sample using the QIAamp PowerFecal Pro DNA extraction kit (Qiagen, Germantown, MD, USA) according to the manufacturer’s instructions. One tube per batch was reserved as a negative control and was treated identically to all samples. Resultant DNA samples were quantified using the Qubit Broad-Range assay kit (Thermo Fisher Scientific, Waltham, MA) on a Synergy 2 Multi-Mode plate reader (BioTek, Winooski, VT).

For metagenomic shotgun sequencing, DNA was submitted to the University of Wisconsin-Madison Biotechnology Center for 2 x 150 bp paired-end sequencing on a Illumina NovaSeq 6000 with a target read depth of 70 million reads per sample, which was deemed sufficient for ARG detection in bovine feces^49^.

### Sequence cleanup and classification

Fastq files from 8 flow cell sequencing lanes were concatenated per sample and resulted in 27.3 billion raw reads from the 116 samples, averaging 235 million reads per sample. These sequences were processed using the AMR++ pipeline v3.0.2^50^ with default settings. In short, sample reads were trimmed using Trimmomatic v0.39^51^, host reads were identified and removed by mapping against the *Bos taurus* reference assembly ARS-UCD1.3, which left 22.7 billion reads. Resultant reads were aligned to the MEGARes database v3.00^50^ using BWA^52^ to identify ARGs. Alignments were deduplicated and genes identified; those that required single nucleotide polymorphism (SNP) confirmation were confirmed using SNP_Verification.py (https://github.com/Isabella136/AmrPlusPlus_SNP, accessed 12/20/2023) with default settings. This ultimately resulted in 11,880,966 deduplicated and SNP-confirmed resistance gene counts, that specifically represented 1,317 ARGs that confer resistance to 25 antimicrobial classes through 84 mechanisms, used for subsequent analyses. Finally, reads that aligned to the MEGARes database were extracted using samtools^53^ and taxonomically classified for each sample using Kraken 2 v2.1.3^54^ with a confidence cutoff of 0.1 against the Kraken 2 PlusPFP index (downloaded 11/29/2023).

### Statistical analysis

Statistical analyses were also performed in R. The deduplicated and SNP-confirmed count dataset was subset to include only “Drug” resistance genes, removing genes conferring resistance to biocides, metals, or multi-compounds. The ARG count matrix was imported into phyloseq v1.44.0^55^ where the alpha diversity metrics, Chao’s Richness and Shannon’s Diversity Index, were then calculated. When evaluating the significance of variable impact on resistome alpha diversity, the Wilcoxon test was used for binary variables and the Kruskal-Wallis (KW) test was used for multinomial variables for this non-parametric data. Post-hoc pairwise comparisons between cattle groups were performed using the Wilcoxon Sum Rank test with FDR-adjusted p-values (*P* < 0.05).

ARG counts were normalized to account for differences in sequencing depth using cumulative sum scaling (CSS) (cumNorm::metagenomeSeq)^56^ as previously described^36^. Sankey plots (sankeypq::MiscMetabar) were created using normalized data for antimicrobial resistance class as the node, antimicrobial resistance mechanisms and antimicrobial resistance genes as links, and cattle groups as the targets. Beta-diversity of drug class was calculated using Bray-Curtis Dissimilarity and visualized using nonmetric multi-dimensional scaling (NMDS) plots with automatic WI double-standard transformation (ordinate::phyloseq)^57^. A stressplot was generated to analyze the non-metric fit (stressplot::vegan). Of note, resistomes were not found to differ across longitudinal sampling for any diversity metric (*P* > 0.05) and thus the longitudinal sampling variable was not included in subsequent models.

Beta-dispersion (PERMDISP) (vegan::betadisper) was used to test data for the homogeneity of variance assumption for permutational multivariate ANOVA (PERMANOVA) (vegan::adonis2) analyses which were used to assess significance as appropriate with permutations = 10,000. When analyzing the impact of AMU (low or high), farm was included as a random effect variable and cattle group was included as strata to restrict the permutations within each group. For analysis of cattle groups, AMU (low or high) was included as a fixed effect variable and farm was included as a random effect variable.

Differential abundance of resistance classes was determined for pairwise comparison of cattle groups using unnormalized data due to the automatic normalization during Analysis of Compositions of Microbiomes with Bias Correction 2 (ANCOMBC::ancombc2). Farm was included again as a random effect variable, with AMU and cattle group as fixed effects. Sensitivity analysis was performed to ensure the robustness of differential abundance results against differing pseudo-counts being added to zeros. Mixed directional false discovery rate (mdFDR) was used to adjust p-values for pairwise comparisons across cattle groups.

Zero inflated gaussian models (metagenomeSeq::fitZig) were used to identify individual resistance genes that were differentially abundant between variable groups. Empirical Bayes moderation (limma::eBayes) was applied to the contrasted fit of the ZIG model in order to account for variance estimates across taxa to provide robust estimates for gene features. Benjamini-Hochberg (BH) false discovery rate (FDR) was used for p-value adjustment. These models were used to create volcano plots that display ARGs with a significant (*P* < 0.05) log fold change |logFC value| > 1.5 between variable groups.

## Author Contributions

Conceptualization, A.K.S, G.S., J.M.S, N.S., T.L.G., and P.L.R; methodology, A.K.S, G.S., J.M.S., N.S., T.L.G., P.L.R., A.J.S. and J.L.d.C.; software, A.J.S., C.A.N.; validation, G.S. and P.L.R.; formal analysis, A.J.S., C.A.N.; investigation, A.J.S. and J.L.d.C.; resources G.S. and P.L.R.; data curation, A.J.S., C.A.N., and J.L.d.C.; writing-original draft preparation, A.J.S., C.A.N., G.S., and J.L.H.; writing-review and editing, A.J.S., C.A.N., G.S., J.L.H., J.L.d.C, N.S., A.E.K., A.K.S., J.M.S., T.L.G., and P.L.R.; visualization, A.J.S., C.A.N., G.S., and J.L.H., supervision, P.L.R; project administration, A.E.K. and P.L.R.; funding acquisition, A.K.S., G.S., J.M.S., N.S., T.L.G., and P.L.R. All authors have read and agreed to the published version of the manuscript.

## Review Board Statement

The animal study protocol was approved by the University of Wisconsin Institutional Animal Care and Use Committee (IACUC) under protocol A005774.

## Data Availability Statement

The sequencing datasets generated and analyzed during the current study are available in the NCBI Sequence Read Archive (SRA) repository under BioProject number PRJNA1093066.

## Acknowledgements

We would like to thank the dairy farms enrolled in this study for their participation. We would also like to thank the Suen lab for their support and invaluable suggestions.

