## Supplemental Information for "Antimicrobial use contributes to resistance gene enrichment across cattle groups on commercial dairy farms"

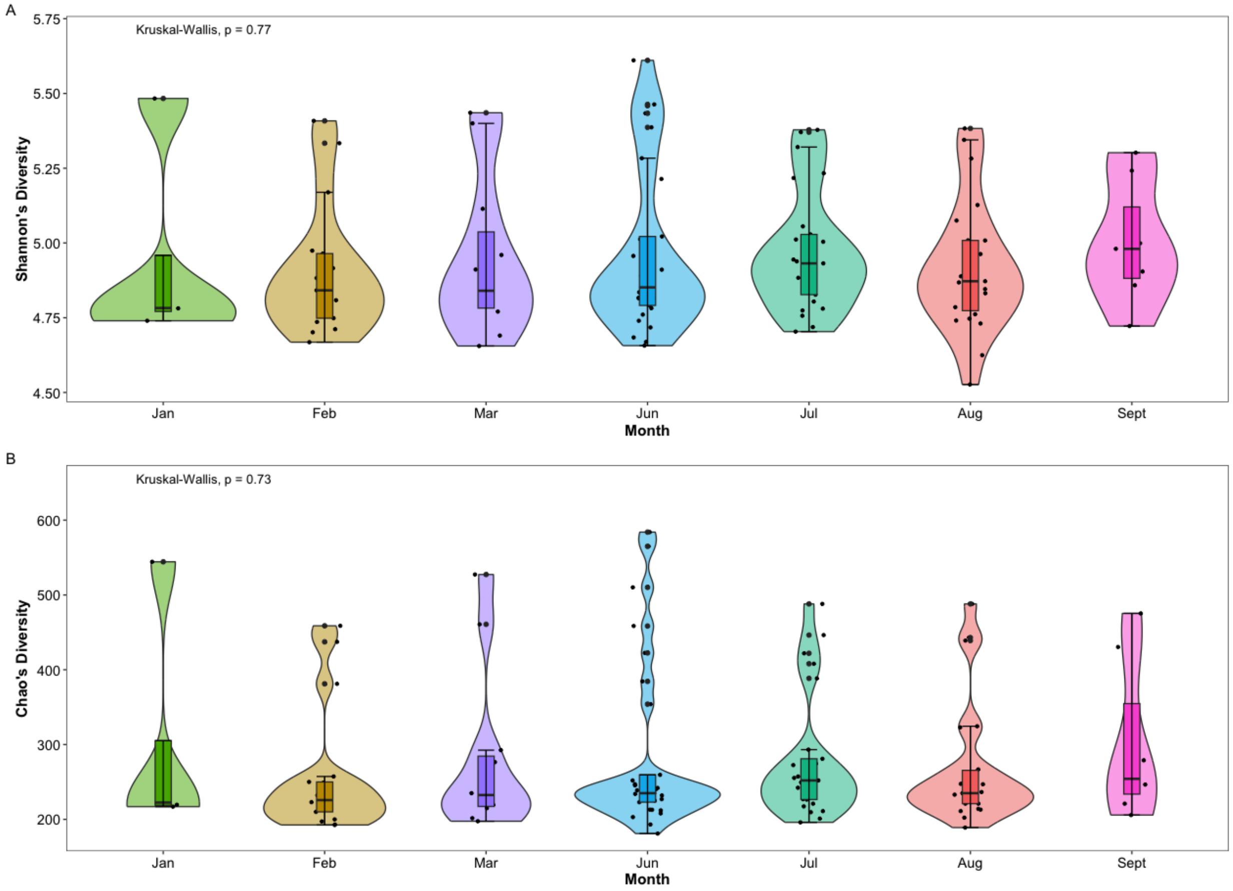


**Supplemental Fig. 1** Impact of month on (A) Shannons's diversity (Kruskal-Wallis Test $P_{FDR\text{-}adj}$ = 0.77) and (B) Chao's richness (Kruskal-Wallis Test $P_{FDR\text{-}adj}$ = 0.73) on cattle resistomes. The Wilcoxon Ranked Sums Test was used to identify significant pairwise comparisons between each season, but none were significant.

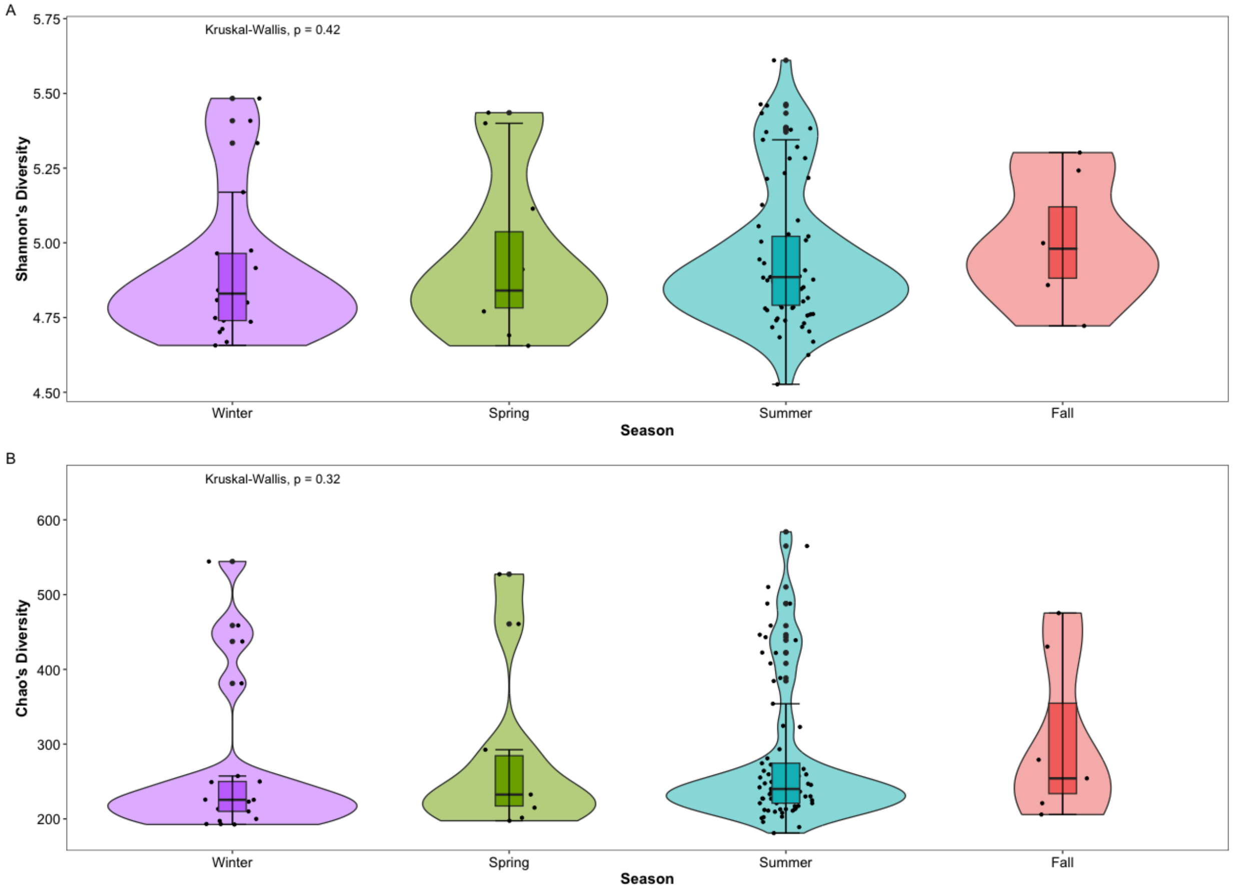


**Supplemental Fig. 2** Impact of season on (A) Shannons's diversity (Kruskal-Wallis Test $P_{FDR\text{-}adj}$ = 0.42) and (B) Chao's richness (Kruskal-Wallis Test $P_{FDR\text{-}adj}$ = 0.32) on cattle resistomes. The Wilcoxon Ranked Sums Test was used to identify significant pairwise comparisons between each season, but none were significant.

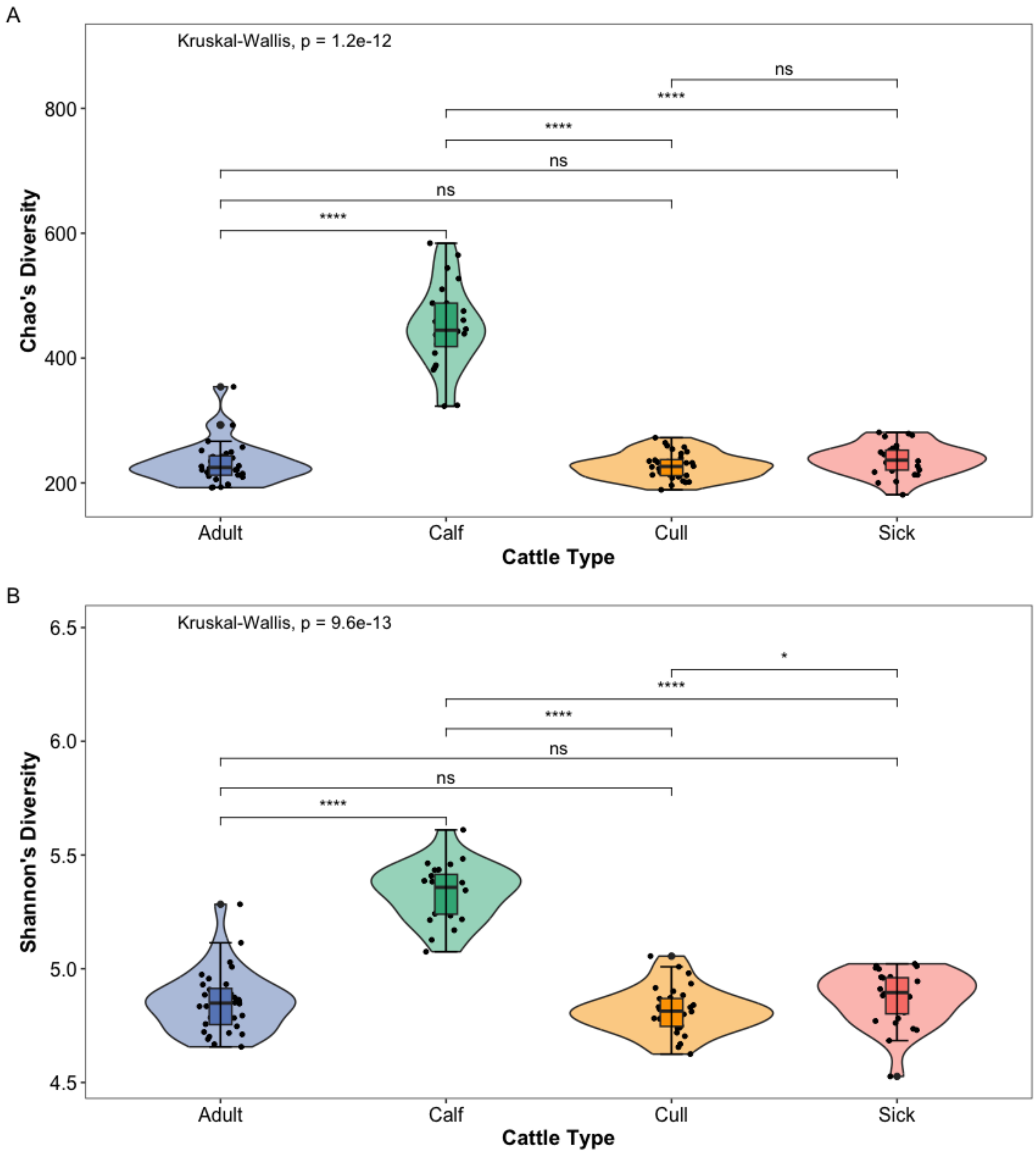


**Supplemental Fig. 3** Impact of cattle group on (A) Chao's richness (Kruskal-Wallis Test $P_{FDR\text{-}adj}$ = 1.2e-12) and (B) Shannons's diversity (Kruskal-Wallis Test $P_{FDR\text{-}adj}$ = 9.6e-13) of the cattle resistomes. The Wilcoxon Ranked Sums Test was used to identify significant pairwise comparisons between each cattle group, indicated by, < 0.05 by *, < 0.001 by ***, and <0.0001 by ****.

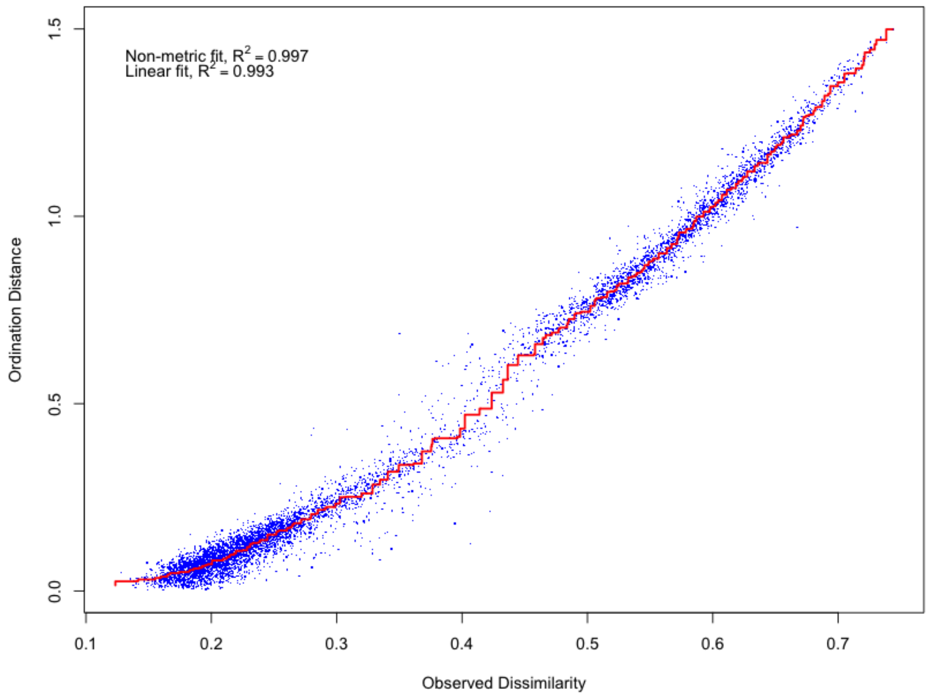


**Supplemental Fig. 4**. Stressplot of Bray-curtis's distance measures for resistome class. Stress = 0.058 indicating a good fit. Non-metric fit $R^2$ = 0.997 also indicates a good fit.

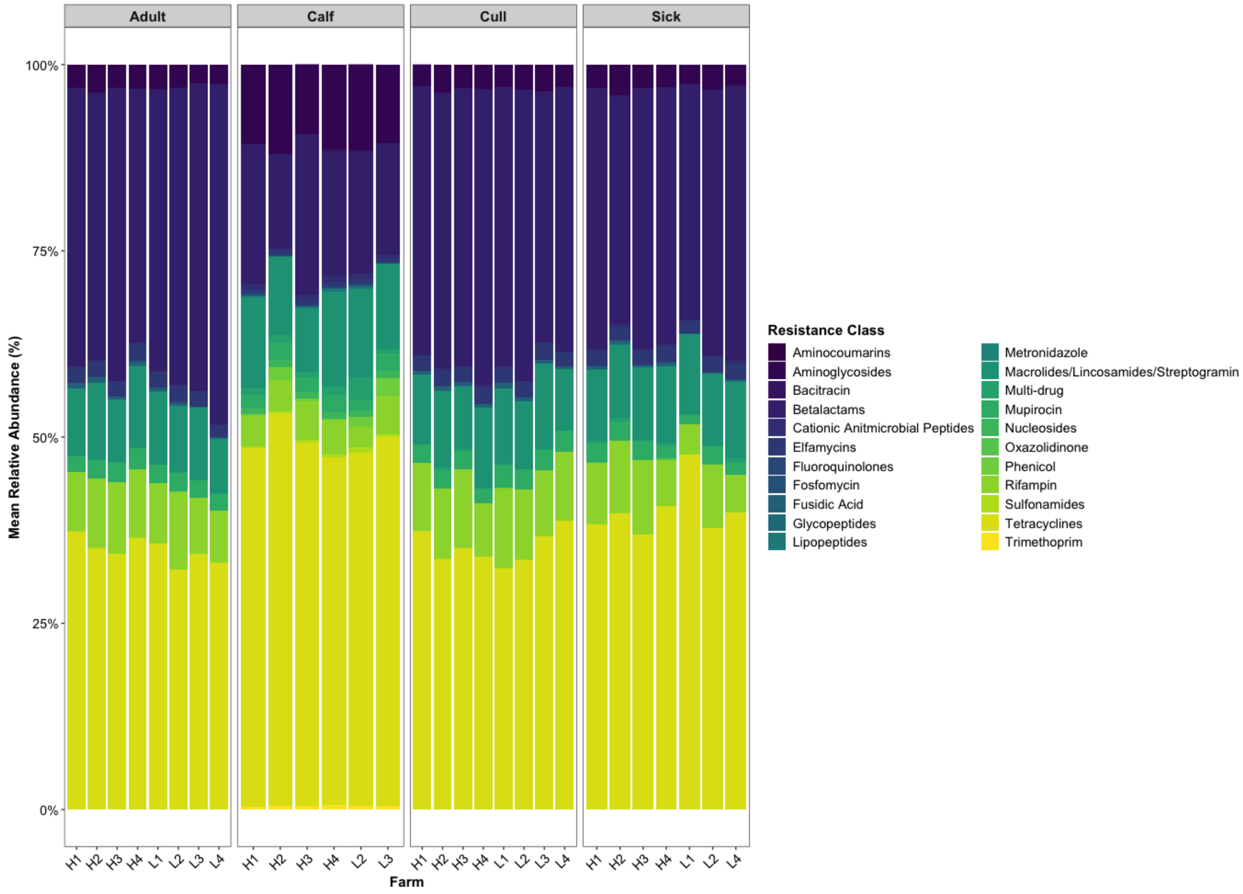


**Supplemental Fig. 5** Relative abundance representing > 0.1% of total gene accession counts across all samples of resistance class composition in cattle at each farm by cattle group.

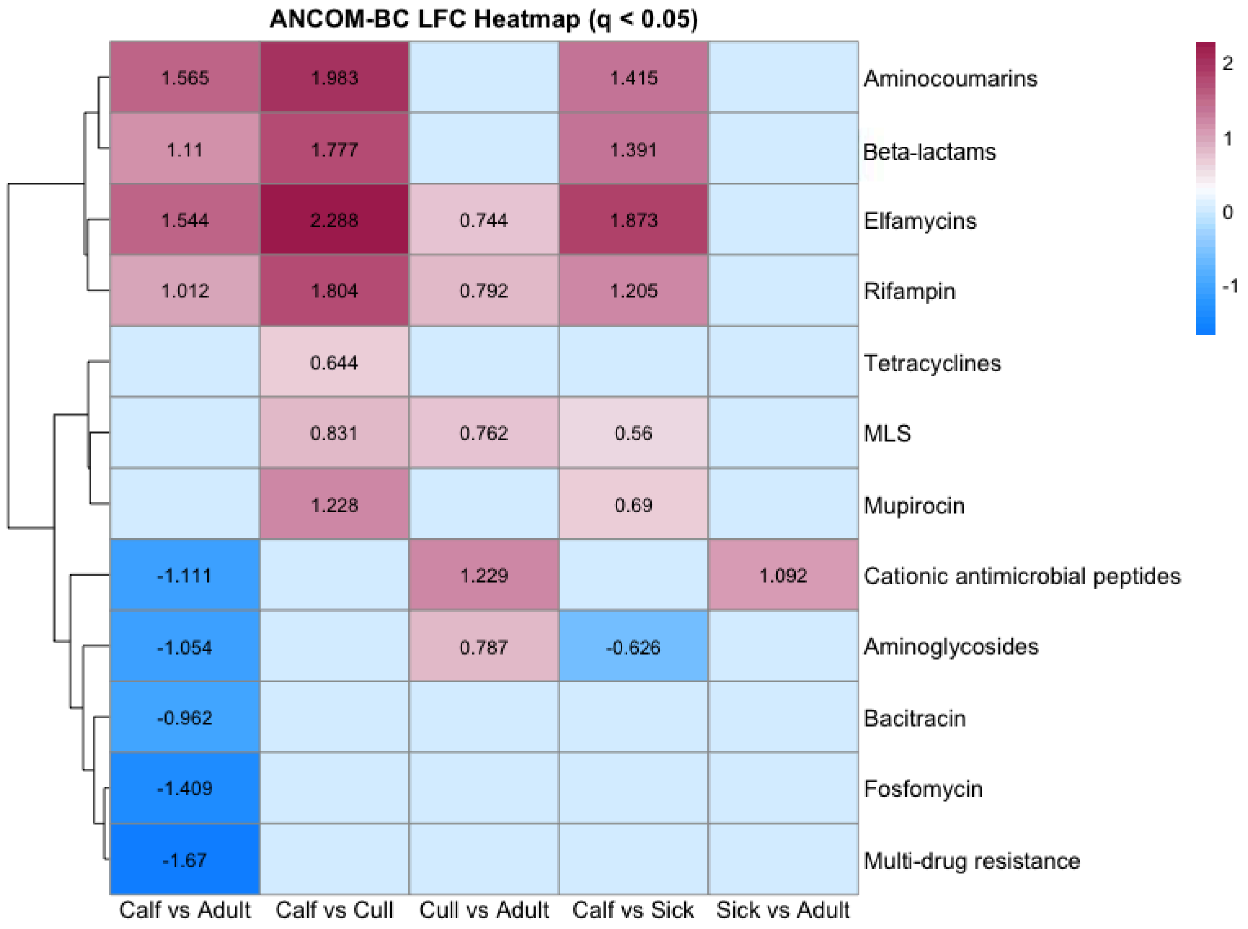


**Supplemental Fig. 6** Differentially abundant antimicrobial resistance classes between cattle groups. Heatmap of differentially abundant antimicrobial resistance classes for each pairwise comparison of calves (Calf), cull cows (Cull), sick cows (Sick), and healthy lactating cows (Adult). Colors represent log fold change in abundance, red for increased log fold change in the right group compared to the left, and blue for decreased log fold change in the right group compared to the left. Comparisons with FDR-corrected p-values > 0.05 were set to 0 and colored white. Aminocoumarins, Beta-lactams, elfmycins, and rifampin were all consistently higher in adult cows compared to calves. Alternatively, cationic antimicrobial peptides, aminoglycosides, bacitracin, fosfomycin, and multi-drug resistance were all reduced in adults compared to calves.

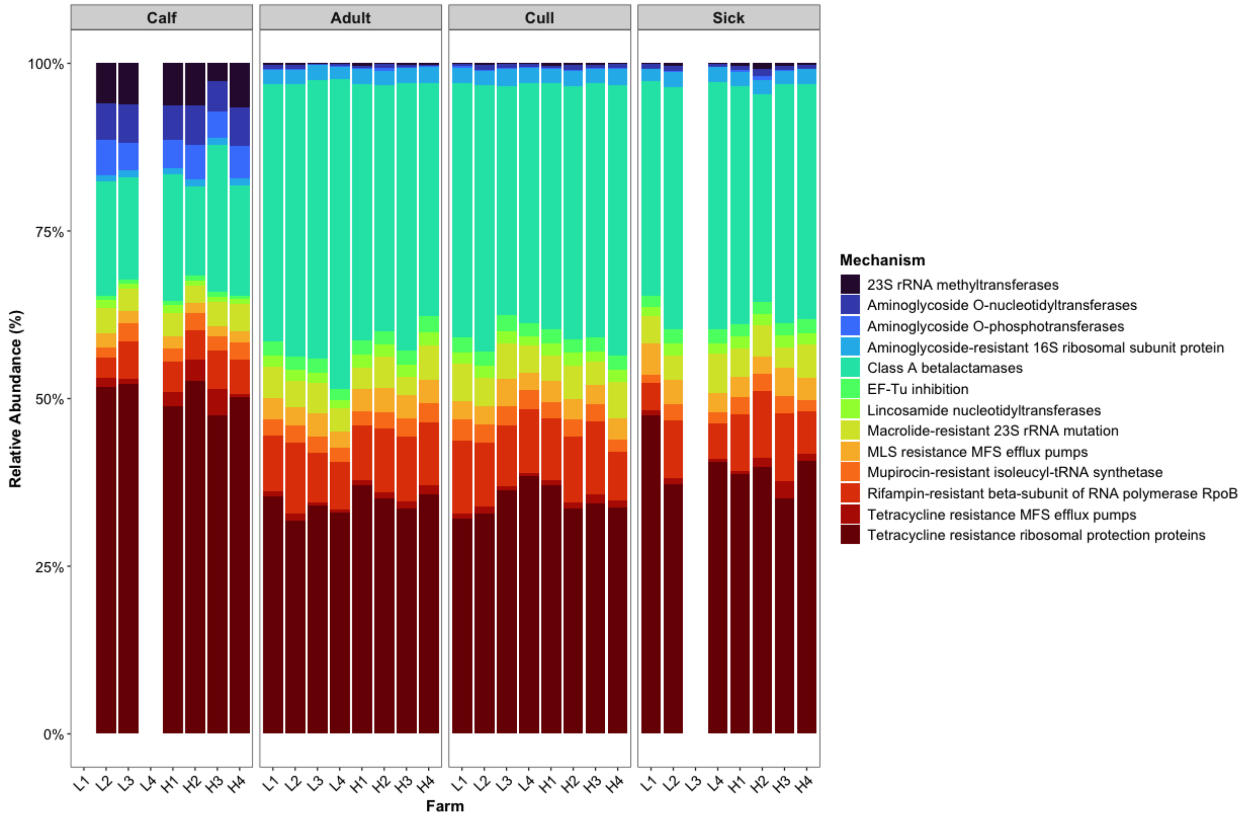


**Supplemental Fig. 7** Relative abundance representing > 0.5% of total counts across all samples for resistance mechanism composition in cattle at each farm by cattle group.

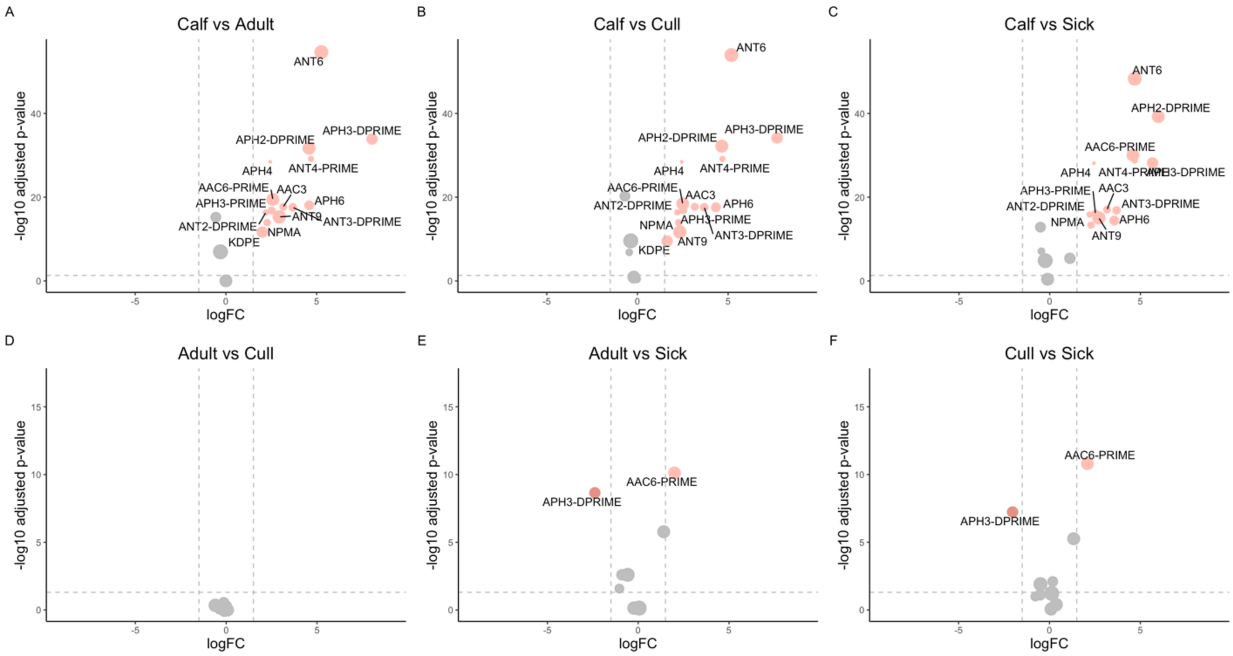


**Supplemental Fig. 8** Zero-inflated gaussian models were used to determine which aminoglycoside resistance genes were significantly (Benjamini-Hochberg False Discovery Rate adjusted p-value < 0.05) higher (right: log Fold Change (logFC) > + 0.5) or lower (left: logFC < -0.5) in abundance between each cattle group. The -log10 adjusted p-value is displayed on the y-axis with the horizontal gray dashed line indicating the significance cut-off (-log10 (0.05)). The logFC in gene abundance between each management system is indicated by circle size and displayed on the x-axis with the vertical dashed gray lines indicating an increased (logFC > + 1.5) or decreased abundance (logFC < -1.5) cutoff.

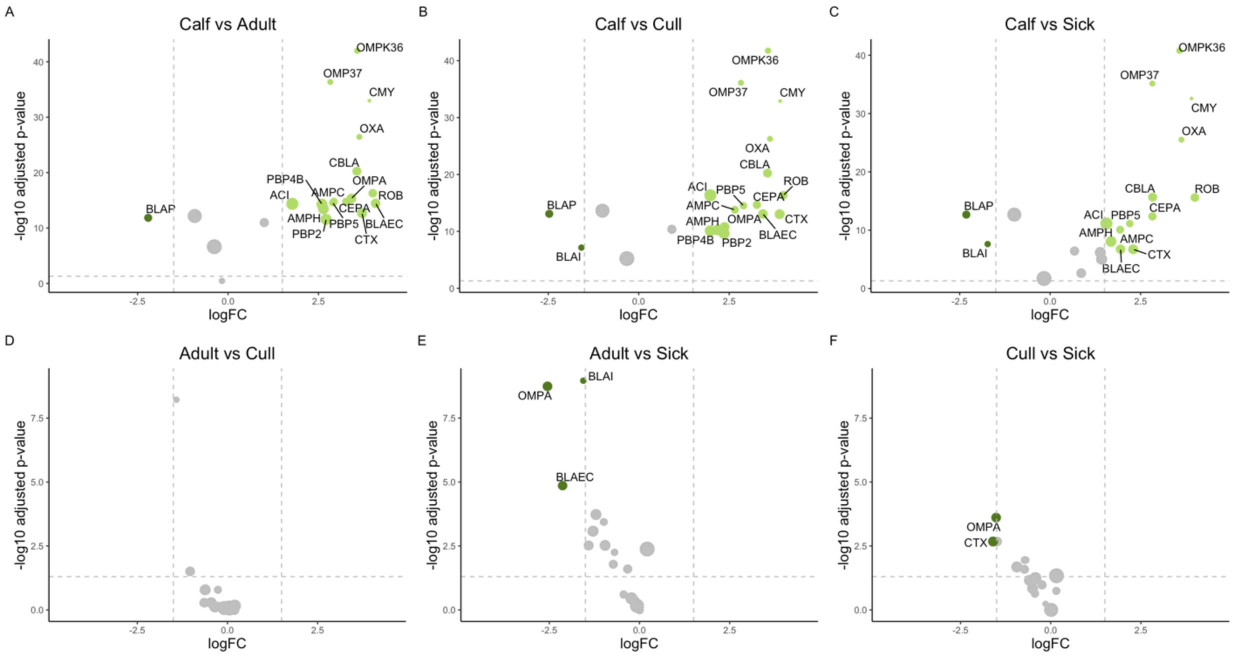


**Supplemental Fig. 9** Zero-inflated gaussian models were used to determine which β-lactam resistance genes were significantly (Benjamini-Hochberg False Discovery Rate adjusted p-value < 0.05) higher (right: log Fold Change (logFC) > + 0.5) or lower (left: logFC < -0.5) in abundance between each cattle group. The -log10 adjusted p-value is displayed on the y-axis with the horizontal gray dashed line indicating the significance cut-off (-log10 (0.05)). The logFC in gene abundance between each management system is indicated by circle size and displayed on the x-axis with the vertical dashed gray lines indicating an increased (logFC > + 1.5) or decreased abundance (logFC < -1.5) cutoff.

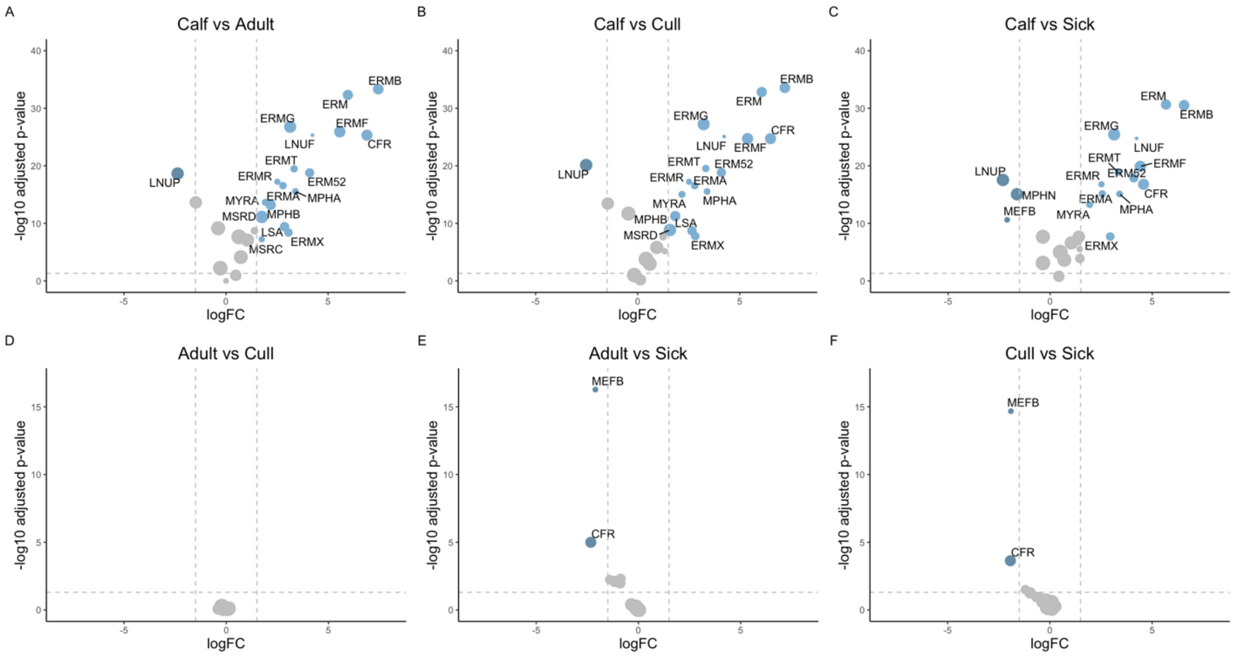


**Supplemental Fig. 10** Zero-inflated gaussian models were used to determine which macrolide/lincosamide/streptogramin resistance genes were significantly (Benjamini-Hochberg False Discovery Rate adjusted p-value < 0.05) higher (right: log Fold Change (logFC) > + 0.5) or lower (left: logFC < -0.5) in abundance between each cattle group. The -log10 adjusted p-value is displayed on the y-axis with the horizontal gray dashed line indicating the significance cut-off (-log10 (0.05)). The logFC in gene abundance between each management system is indicated by circle size and displayed on the x-axis with the vertical dashed gray lines indicating an increased (logFC > + 1.5) or decreased abundance (logFC < -1.5) cutoff.

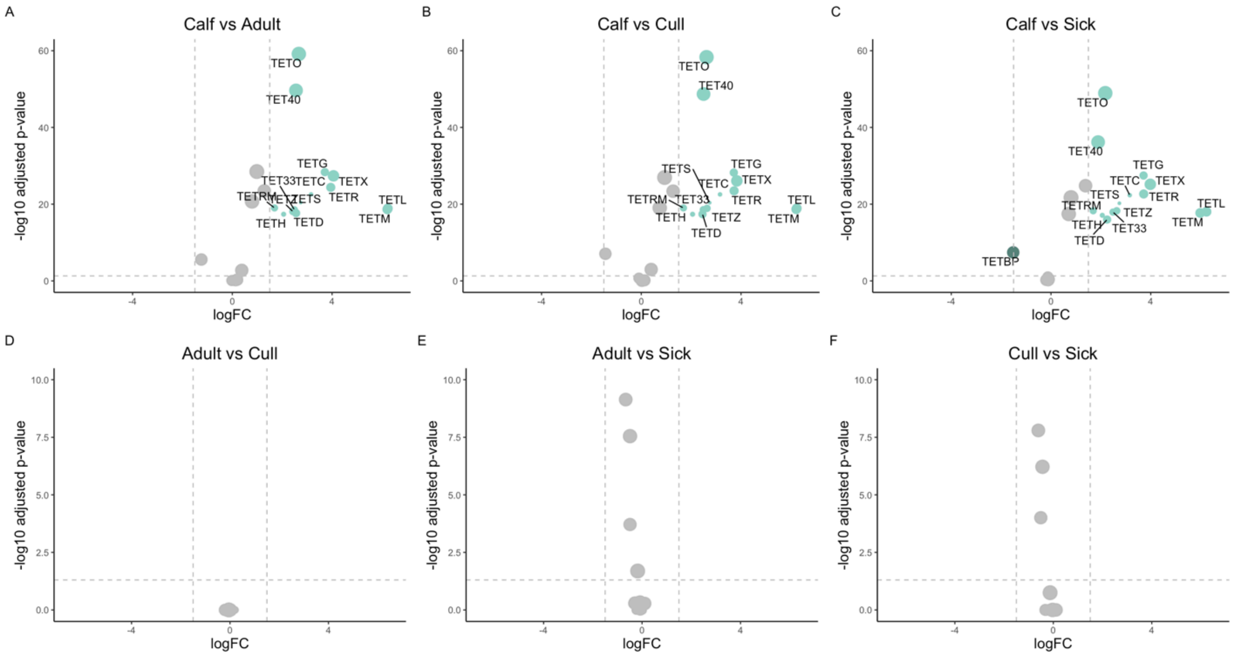


**Supplemental Fig. 11** Zero-inflated gaussian models were used to determine which tetracycline resistance genes were significantly (Benjamini-Hochberg False Discovery Rate adjusted p-value < 0.05) higher (right: log Fold Change (logFC) > + 0.5) or lower (left: logFC < -0.5) in abundance between each cattle group. The -log10 adjusted p-value is displayed on the y-axis with the horizontal gray dashed line indicating the significance cut-off (-log10 (0.05)). The logFC in gene abundance between each management system is indicated by circle size and displayed on the x-axis with the vertical dashed gray lines indicating an increased (logFC > + 1.5) or decreased abundance (logFC < -1.5) cutoff.

**Supplemental Table 1.** PERMANOVA pairwise comparisons of resistance class variance by antimicrobial use (AMU) level. PERMDISP homogeneity of groups was tested to ensure PERMANOVA assumptions were met.

| **Variance of specific resistance classes by AMU level** | | | |
|---|---|---|---|
| | **PERMANOVA $R^2$** | **PERMANOVA *P-val*** | **PERMDISP *P-val*** |
| **Resistance Classes** | 0.086 | 0.003 | 0.940 |
| **Aminoglycosides** | 0.085 | 0.028 | 0.620 |
| **Betalactams** | 0.128 | 0.015 | 0.305 |
| **Elfamycins** | 0.022 | 0.997 | 0.495 |
| **MLS** | 0.129 | 0.0003 | 0.840 |
| **Mupirocin** | 0.073 | 0.286 | 0.930 |
| **Tetracyclines** | 0.057 | 0.758 | 0.810 |
| **Rifampin** | 0.105 | 0.045 | 0.100 |
| **Variance of all genes within specific resistance Genes by AMU level** | | | |

| | PERMANOVA $R^2$ | PERMANOVA *P-val* | PERMDISP *P-val* |
|---|---|---|---|
| **Resistant Genes** | 0.081 | 0.0341 | 0.985 |
| **Aminoglycoside Genes** | 0.083 | 0.018 | 0.685 |
| **Betalactam Genes** | 0.110 | 0.013 | 0.560 |
| **Elfamycin Genes** | 0.022 | 0.998 | 0.565 |
| **MLS Genes** | 0.144 | 0.00001 | 0.950 |
| **Mupirocin Genes** | 0.073 | 0.284 | 0.495 |
| **Tetracycline Genes** | 0.070 | 0.288 | 0.630 |
| **Rifampin Genes** | 0.095 | 0.072 | 0.865 |

**Supplemental Table 2.** Richness and diversity pairwise comparisons p-values in regards to

| **Chao** | | Adult | Calf | Cull |
|---|---|---|---|---|
| | Calf | <0.0001 | - | - |
| | Cull | ns | <0.0001 | - |
| | Sick | ns | <0.0001 | ns |
| **Shannon** | | | | |
| | Calf | <0.0001 | - | - |
| | Cull | ns | <0.0001 | - |
| | Sick | ns | <0.0001 | <0.05 |

**Supplemental Table 3.** PERMANOVA for the beta diversity of resistant genes in specific resistance classes by cattle group

| | PERMANOVA | PERMDISP |
|---|---|---|
| **Resistance Class** | <0.001 | 0.022 - fail |
| **Betalactam Class** | <0.001 | 0.099 - pass |
| **Elfamycin Class** | <0.001 | 0.002 - fail |
| **Tetracycline Class** | <0.001 | 0.550 - pass |
| **Macrolides Class** | <0.001 | 0.156 - pass |
| **Mupirocin Class** | <0.001 | 0.005 - fail |
| **Aminoglycoside Class** | <0.001 | 0.554 - pass |
| **Rifampin Class** | 0.002 | 0.004 - fail |

**Supplemental Table 4.** PERMANOVA for the pairwise comparisons of resistance class beta diversity by cattle group.

| | ***r2*** | ***P - value*** |
|---|---|---|
| **All AMU Levels** | | |
| **Adult vs Cull** | 0.053 | 0.118 |

| | | |
|---|---|---|
| **Adult vs Sick** | 0.208 | 0.001 |
| **Cull vs Sick** | 0.181 | 0.001 |
| **High AMU** | | |
| *Adult vs Cull* | 0.203 | 0.079 |
| *Adult vs Sick* | 0.296 | 0.001 |
| *Cull vs Sick* | 0.215 | 0.038 |
| **Low AMU** | | |
| *Adult vs Cull* | 0.261 | 0.017 |
| *Adult vs Sick* | 0.465 | 0.001 |
| *Cull vs Sick* | 0.446 | 0.001 |

**Supplemental Table 5.** Spearman's rank correlation values for AMU (animal/DDD) and Chao's Richness or AMU (animal/DDD) and Shannons's Diversity.

| | ***r*** | ***P - value*** |
|---|---|---|
| **Chao's Richness** | | |
| **All Cattle** | 0.223 | 0.016 |
| **Calves** | 0.058 | 0.787 |
| **All Adult Cows** | 0.242 | 0.020 |
| *Healthy Adult Cows* | 0.018 | 0.920 |
| *Cull Adult Cows* | 0.406 | 0.021 |
| *Sick Adult Cows* | 0.369 | 0.053 |
| **Shannon's Diversity** | | |
| **All Cattle** | 0.164 | 0.078 |
| **Calves** | 0.148 | 0.490 |
| **All Adult Cows** | 0.142 | 0.176 |
| *Healthy Adult Cows* | 0.043 | 0.816 |
| *Cull Adult Cows* | 0.388 | 0.028 |
| *Sick Adult Cows* | 0.093 | 0.638 |
